# Structural and mechanistic analysis of covalent ligands targeting the RNA-binding protein NONO

**DOI:** 10.1101/2025.06.10.658857

**Authors:** Garrett L. Lindsey, Thomas K. Hockley, Alejandro Villa Gomez, Andew C. Marshall, William R. Brothers, Colin T. Finney, Jacob Gross, Archa H. Fox, Gene W. Yeo, Bruno Melillo, Charles S. Bond, Benjamin F. Cravatt

**Author notes:** These authors contributed equally to the manuscript (Thomas K. Hockley, Alejandro Villa Gomez).

## Abstract

RNA-binding proteins (RBPs) play important roles in mRNA transcription, processing, and translation. Chemical tools are lacking for RBPs, which has hindered efforts to perturb and understand RBP function in cells. We recently reported a chloroacetamide compound (*R*)-SKBG-1 that covalently binds the RBP NONO and stabilizes its interactions with mRNAs, leading to transcriptional remodeling and the suppression of cancer cell growth. Here, we report the crystal structure of an (*R*)-SKBG-1:NONO complex, which confirms covalent modification of cysteine-145 at a pocket proximal to the RNA-binding interface of the protein. We show that this pocket can also be targeted by a lower reactivity chlorofluoroacetamide analog (*R, R*)-GL-373, which retains the pharmacological properties of (*R*)-SKBG-1, including blockade of estrogen receptor expression in breast cancer cells, while displaying much greater proteome-wide selectivity. Our findings thus show that NONO can be targeted by covalent ligands with high specificity to pharmacologically suppress pro-tumorigenic gene products in cancer cells.

**Highlights:**

- Co-crystal structure reveals how covalent ligands stereoselectively bind NONO
- Chlorofluoroacetamides (CFAs) bind NONO with high proteomic selectivity
- CFA ligands remodel cancer cell transcriptomes in a NONO-dependent manner
- CFA ligands suppress cancer cell growth in a NONO-dependent manner

## Introduction

The human genome encodes a broad array of RNA-binding proteins (RBPs) that regulate the processing, maturation, and function of diverse RNA components of the cell. Major roles for RBPs include the splicing,^1^ post-transcriptional modification,^2^ and translational control of messenger RNAs (mRNAs),^3^ the folding and assembly of large ribosomal RNA (rRNA)-protein complexes,^4^ and the processing of small nuclear RNAs (snRNAs).^5^

Genetic mutations in RBPs cause many human diseases, underscoring the importance of this protein class for supporting cellular and organismal physiology.^6,7^ RBPs also have the potential to serve as drug targets for restoring human health through, for instance, the post-transcriptional regulation of pathogenic or beneficial gene products. In a compelling proof-of-principle, the small-molecule drug risdiplam stabilizes binding of the U1 small nuclear ribonucleoprotein (snRNP) complex to weak 5’ splice sites to promote exon inclusion and expression of the *SMN2* gene to treat spinal muscular atrophy (SMA) caused by deleterious mutations in the related *SMN1* gene.^8^ Nonetheless, realizing the broader therapeutic potential of RBPs depends on the identification of chemical probes for additional members of this large protein class, a goal that has been hindered by the limited availability of functional assays for RBPs^9,10^ and their perceived poor general druggability.^11,12^

We recently discovered by integrated phenotypic screening and chemical proteomics a series of electrophilic α-chloroacetamide (CA) compounds that covalently bind the RBP NONO to transcriptionally remodel and impair the growth of human cancer cells.^13^ NONO is a member of the Drosophila Behaviour/Human Splicing (DBHS) family of RBPs that includes the paralogs PSPC1 and SFPQ.^14,15^ These abundant nuclear proteins form hetero- and homo-dimers to regulate various stages of mRNA processing, transport, maturation, and stability.^14,15^ The lead compound (*R*)-SKBG-1 was found to engage a paralog-restricted cysteine (C145) located in a hinge region between the RRM1 and RRM2 RNA-binding domains of NONO^16–18^ and mechanistic studies supported a model where (*R*)-SKBG-1 stabilizes NONO binding to mRNAs, including transcripts encoding the androgen receptor and its major splice variants, leading to decreases in their expression in cancer cells.^13^ Consistent with a gain-of-toxic function mechanism that cannot be overcome by the compensatory action of PSPC1 and SFPQ, the pharmacological effects of (*R*)-SKBG-1 were abrogated in cancer cells genetically disrupted for NONO.^13^

While (*R*)-SKBG-1 has provided a valuable initial tool compound for studying NONO function in cells, our understanding of the structural basis for (*R*)-SKGB-1-NONO interactions remains limited. Additionally, (*R*)-SKBG-1 covalently modified many other proteins in cancer cells and showed only a narrow window of stereoselective antiproliferative effects compared to the inactive enantiomeric control compound (*S*)-SKBG-1. These deficiencies likely relate, at least in part, to the high intrinsic reactivity of the CA reactive group of (*R*)-SKBG-1. Here, we report the X-ray crystal structure of a (*R*)-SKBG-1:NONO complex, which revealed a pocket supporting stereoselective reactivity at C145 that is proximal to the predicted RNA-binding interface of NONO. We concurrently discover that NONO can be targeted by an analog of (*R*)-SKBG-1 bearing a lower reactivity chlorofluoroacetamide (CFA) group. This CFA analog (*R, R*)-GL-373 displays stereoselective and NONO-dependent transcriptional remodeling and anti-proliferative activity, while exhibiting much greater proteome-wide selectivity compared to (*R*)-SKBG-1. Among the transcripts suppressed by (*R, R*)-GL-373 in breast cancer cells was the estrogen receptor (ESR1) mRNA, and we verified near complete loss of ESR1 protein by 24 h post-treatment with (*R, R*)-GL-373. Our findings, taken together, support the presence of a druggable and functional pocket in NONO that can be targeted by low-reactivity electrophilic compounds to control pro-tumorigenic transcriptional pathways in cancer cells.

## Results

### (*R*)-SKBG-1-NONO co-crystal structure

We expressed and purified the structured regions of the human NONO homodimer (residues 53-312), NONO/SFPQ heterodimer, and SFPQ homodimer from *E. coli* as previously reported.^19–21^ We then treated purified NONO homodimeric protein (125 µM) with (*R*)-SKBG-1 (4-fold molar excess, 15 min) prior to crystallization. The X-ray crystal structure of the (*R*)-SKBG-1:NONO complex was solved at 2.5 Å resolution and has two NONO homodimers in the asymmetric unit (**Figure 1A** and **Table S1**). Clear electron density was observed for the (*R*)-SKBG-1 compound with a covalent adduct to C145 (in *m*F_o_–*D*F_c_, 3.0 σ omit maps) in each of the four protomers. The sites in chain A and chain B allowed building of the complete (*R*)-SKBG-1 adduct, with chain A having the best-defined electron density, albeit with evidence of dynamic disorder around the sulfonamide-linked methoxyphenyl group (**Figure S1A**). The structure provided a rationale for the stereoselective reactivity of (*R*)-SKBG-1 with C145, as there is a hydrogen bond between the backbone nitrogen of this residue and the carbonyl group adjacent to the (*R*)-SKBG-1 stereocenter (2.9 Å; **Figure 1B**). This hydrogen bond interaction would not be possible with the *(S)*-SKBG-1 enantiomer. Additional contacts that may contribute to orienting (*R*)-SKBG-1 for reaction include a hydrogen bond of the acetamide with Q229 (3.4 Å) and stacking of the phenoxy substituent with the guanidinium moiety of R75 (**Figure 1B**). The previously reported apo-NONO homodimer structure (PDB 5IFM)^18^ showed structural variability across its six homodimers (RMSD 0.6 Å), which is elevated in the (*R*)-SKBG-1:NONO complex (RMSD 0.8-1.1 Å) due to subtle displacements of helices and loops in the coiled-coil and NOPS domains. These differences are, however, within the range observed for different crystal packing environments of the respective structures (**Figure S1B**).

**Figure 1.**
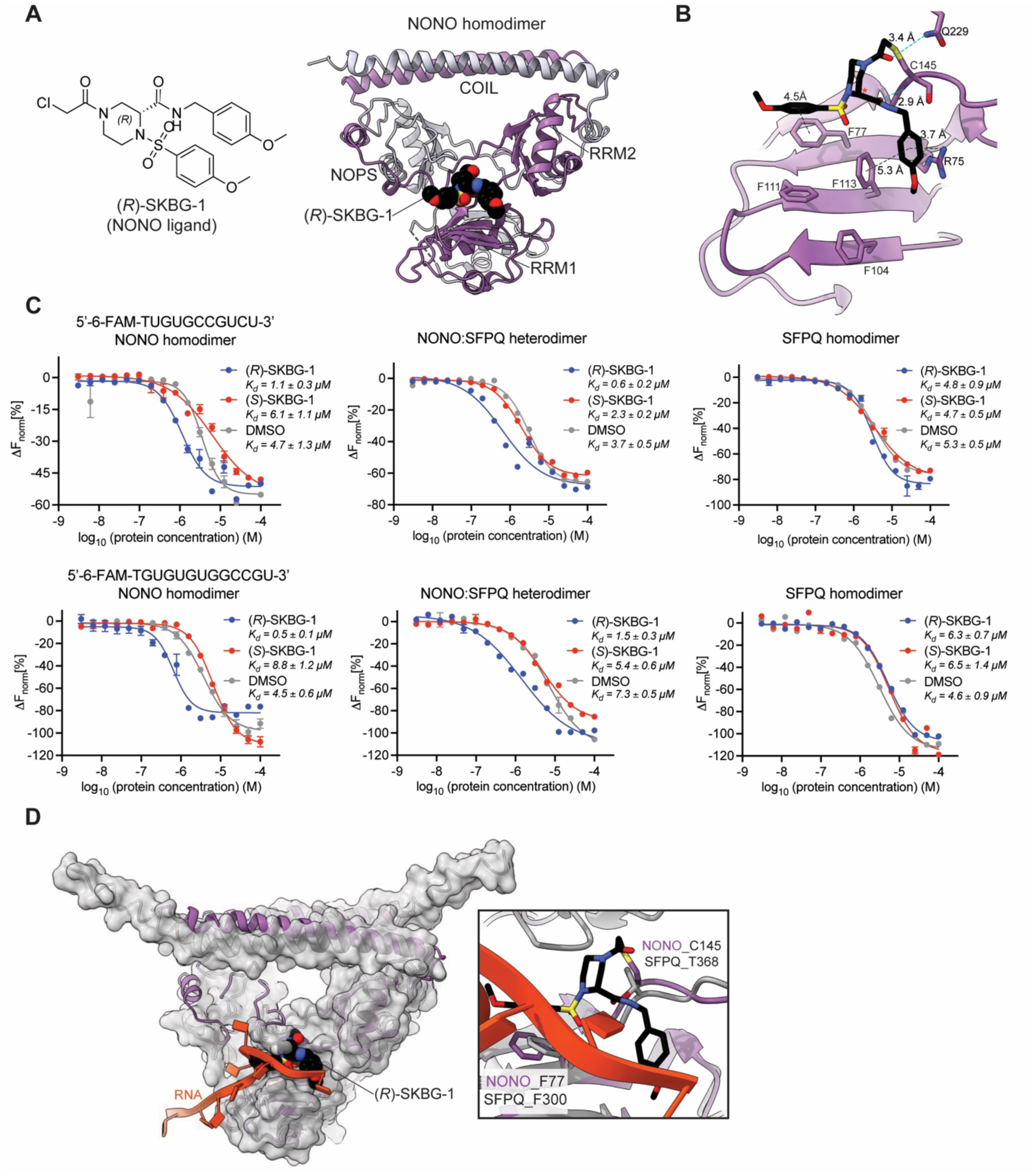
Structural characterization of (*R*)-SKBG-1-bound-NONO homodimer. (A) Left, structure of (*R*)-SKBG-1. Right, 2.5 Å co-crystal structure of (*R*)-SKBG-1-bound NONO homodimer (PDB: 9NZI), where purple and gray mark the individual NONO subunits and (*R*)-SKBG-1 is shown in colored CPK spheres. The RNA-binding domains RRM1 and RRM2 are marked, as are the conserved NOPS and COIL domains. (B) Ligand-binding pocket of NONO highlighting interactions between (*R*)-SKBG-1 and amino acid residues of NONO. The stereocenter in (*R*)-SKBG-1 is marked with a red asterisk (*), as are hydrogen bonds (cyan) from the adjacent carbonyl group to the backbone nitrogen of C145 (2.9 Å) and from the acetamide carbonyl to Q229 (3.4 Å). Interactions between phenyl groups in (*R*)-SKBG-1 and selected sidechains, including conserved phenylalanine residues implicated in RNA interactions (e.g., F77 (4.5 Å), F113 (5.3 Å)) and R75 (3.7 Å), are shown in black. (C) Microscale thermophoresis data for two FAM-labeled RNA oligonucleotides interacting with a NONO homodimer, NONO-SFPQ heterodimer, or SFPQ homodimer in the presence of increasing concentrations of (*R*)-SKBG-1 or (*S*)-SKBG-1. Only the NONO proteins, but not the SFPQ homodimer exhibit a shift to higher RNA binding affinity in the presence of (*R*)-SKBG1 (blue circles). (*S*)-SKBG-1 (red circles) did not alter the RNA binding affinity of the tested proteins in comparison to untreated protein (grey circles). Data are average values ± SD for three independent experiments. (D) Superposition of the (*R*)-SKBG-1-NONO homodimer co-crystal structure (NONO: purple, ligand: black) and a co-crystal structure of SFPQ homodimer bound to RNA (PDB: 7UJ1; SFPQ: gray cartoon and surface, RNA: orange). The inset highlights where (*R*)-SKBG-1 and RNA independently stack with a conserved phenylalanine residue in NONO (F77) and SFPQ (F300), respectively. T368 is the residue in SFPQ that corresponds to C145 in NONO.

The pendant aromatic groups of (*R*)-SKBG-1 are proximal to the sidechains of conserved phenylalanine residues (F77, F113) in the RRM1 domain of NONO that are implicated in RNA-binding^22,23^ (**Figure 1B**), thus providing a potential basis for the compound’s impact on NONO-RNA interactions. We previously found using enhanced UV crosslinking and immunoprecipitation followed by sequencing (eCLIP–seq)^24^ that (*R*)-SKBG-1 stabilizes NONO binding to mRNAs in cells. Here, we evaluated the impact of (*R*)-SKBG-1 on purified NONO interactions with two GU-rich RNA oligonucleotides by microscale thermophoresis. These experiments revealed that (*R*)-SKBG-1, but not the inactive enantiomer (*S*)-SKBG-1, increased the affinity of the NONO homodimer or NONO:SFPQ heterodimer for single-stranded RNA (**Figure 1C** and **Figure S1C, D**). In contrast, (*R*)-SKBG-1 did not alter the RNA affinity of an SFPQ homodimer (**Figure 1C**).

Finally, alignment of the (*R*)-SKBG-1:NONO complex structure with an RNA-bound structure of the SFPQ homodimer (PDB: 7UJ1)^22^ revealed that (*R*)-SKBG-1 occupies a site overlapping with RNA in the SFPQ structure (**Figure 1D**). Considering that our biochemical and cell biological studies indicate (*R*)-SKBG-1 stabilizes rather than disrupts NONO-RNA interactions, we speculate that the (*R*)-SKBG-1-bound NONO protein may adopt different modes of RNA interaction that avoid steric clashes with the covalent (*R*)-SKBG-1:C145 adduct. The observations from prior studies that the RNA-binding interface of RRM1 is quite open and flat, and that the RRM1 domains of both NONO and SFPQ have been shown to move to produce a more closed state on binding nucleic acid^18,22^ are consistent with this hypothesis.

### Chlorofluoroacetamides engage NONO with improved proteome-wide selectivity

While original CA ligands such as (*R*)-SKBG-1 engage NONO_C145 with excellent stereoselectivity, the broader proteomic reactivity of these compounds is high,^13^ which limits their utility in some types of cell biology experiments. In considering alternative approaches to covalently engage NONO with less reactive electrophiles, we noted that the hydrogen bond between the backbone N-H of C145 and the C2-carboxamide carbonyl of (*R*)-SKBG-1, and between the sidechain of Q229 and the CA carbonyl of (*R*)-SKBG-1 (**Figure 1B**), together positioned the electrophilic α-carbon center of this compound for reactivity with C145. We accordingly inferred that substitution of the CA group with lower reactivity β-carbon electrophiles like an acrylamide or butynamide might create compounds that are misaligned for covalent engagement of NONO_C145. As an alternative, we considered analogs bearing a chlorofluoroacetamide (CFA), which closely mimics the CA geometry while exhibiting much lower electrophilicity and has emerged as an attractive reactive group for the design of covalent ligands with improved selectivity.^25–27^

A full set of CFA stereoisomeric analogs of (*R*)- and (*S*)-SKBG-1 was synthesized (**Figure 2A**) and evaluated for reactivity with NONO by cysteine-directed activity-based protein profiling (ABPP)^28–31^ in the prostate cancer cell line 22Rv1 (20 µM compound, 6 h). Of the four CFA analogs, a single stereoisomer (*R, R*)-GL-373 showed robust engagement of NONO_C145 (**Figure 2B** and **Dataset S1**). The inactivity of (*S, R*)-GL-373 indicated that the stereoconfiguration at the CFA center was crucial for reactivity with NONO_C145, as has been observed for CFA ligands targeting other proteins (e.g., SARS-CoV2 protease^26^). We next generated alkyne analogs of (*R, R*)- and (*S, S*)-GL-373 – (*R, R*)- and (*S, S*)-GL-586, respectively – and evaluated the reactivity of these compounds by gel-ABPP in 22Rv1 cells stably expressing WT-NONO or a C145S-NONO mutant. These experiments showed clear stereoselective and site-specific reactivity of (*R, R*)-586 with WT-NONO with only a handful of additional protein reactivity events being observed across the gel profile (**Figure 2C**). The (*R, R*)-GL-586-WT-NONO interaction was blocked by pre-treatment with (*R, R*)-GL-373 (20 µM, 6 h), but not other CFA stereoisomers (**Figure 2D**). Using this competitive gel-ABPP assay, we determined half-maximal target engagement (TE_50_) values for NONO of 2.2 and 9.0 µM for (*R*)-SKBG-1 and (*R, R*)-GL-373, respectively (**Figure 2E**). We also noted that (*R, R*)-GL-373 produced a partial stereoselective blockade of a strong ∼55 kDa signal (**Figure 2D**) found in both WT-NONO or a C145S-NONO-expressing 22Rv1 cells (**Figure 2C**), as well as in cells genetically disrupted for NONO (sgNONO cells; **Figure S2**). We speculate that this signal corresponds to a combination of endogenous NONO and an additional target of the (*R, R*)-GL-586 probe.

**Figure 2.**
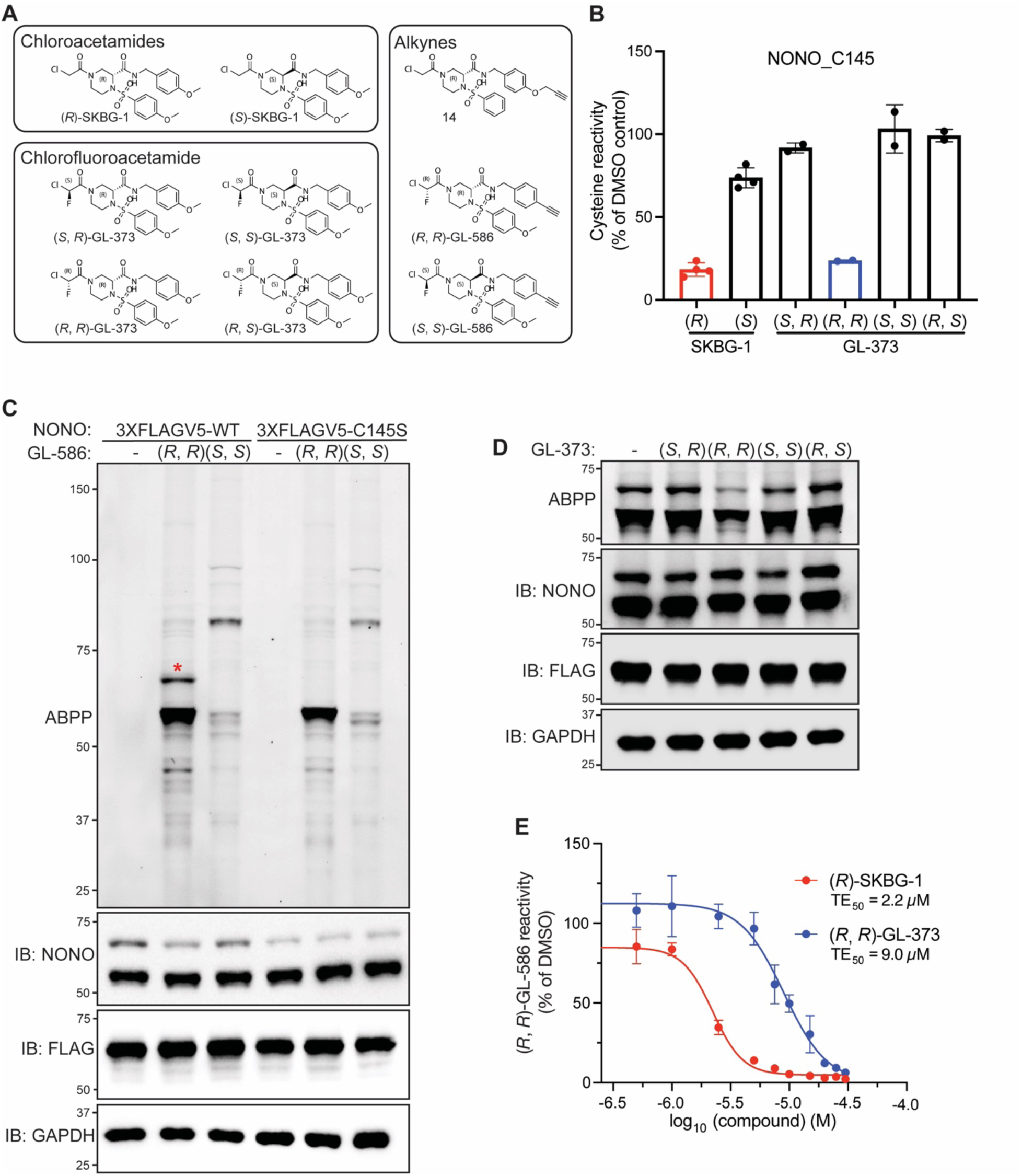
Chlorofluoroacetamide (CFA) ligands stereoselectively and site-specifically engage NONO_C145. (A) Structures of chloroacetamide (CA) and chlorofluoroacetamide (CFA) compounds used in this study. Left boxes show parent compounds and right box shows corresponding alkyne analogs. (B) Bar graph presenting cysteine-directed ABPP data for NONO_C145 from 22Rv1 cells treated with the indicated CA and CFA compounds (20 µM, 6 h). Data are average values ± SD for two-four independent experiments. (C) Gel-ABPP data showing the proteomic reactivity of alkynes (*R, R*)-GL-586 and (*S, S*)-GL-586 (10 *µ*M, 1 h) in 22Rv1 cells stably expressing 3X-FLAG-V5 epitope-tagged WT-NONO [WT-3XFLAGV5] or C145S-NONO [C145S-3XFLAGV5]). GL-586-reactive proteins were detected by conjugation of an azide-rhodamine reporter group using copper-catalyzed azide-alkyne cycloaddition (CuAAC) chemistry,^42,43^ followed by SDS–PAGE, and in-gel fluorescence scanning.^44^ Red asterisk (*) highlights signal corresponding to molecular weight of 3XFLAGV5-NONO. Data are from a single experiment representative of two independent experiments. (D) Gel-ABPP data showing stereoselective blockade of alkyne (*R, R*)-GL-586 reactivity with 3XFLAGV5-WT-NONO in 22Rv1 cells by (*R, R*)-GL-373. Cells were pre-treated with GL-373 stereoisomers (20 µM, 6 h) followed by (*R, R*)-GL-586 (10 µM, 1 h) and gel-ABPP analysis as described in panel (C). (E) Concentration-dependent blockade of (*R, R*)-GL-586 reactivity with WT-NONO by (*R*)-SKBG-1 and (*R, R*)-GL-373 as determined by gel-ABPP. Experiments were performed as described in panel (D), and data are average values ± SD for three independent experiments. TE_50_ values: (*R*)-SKBG-1, 2.2 *µ*M (95% confidence intervals (CI) 2.0–2.5 µM); (*R, R*)-GL-373, 9.0 µM (95% CI 7.6–12 µM).

Despite a modest (∼four-fold) reduction in potency, the CFA ligands displayed much better proteome-wide selectivity compared to CA ligands as determined by gel-ABPP experiments performed with (*R, R*)-GL-586 versus an alkyne analog of (*R*)-SKBG-1 (compound **14**;^13^ **Figure 3A**) and cysteine-directed ABPP experiments assessing the global reactivity of (*R, R*)-GL-373 and (*R*)-SKBG-1 (**Figure 3B** and **Dataset S1**). In the gel-ABPP experiments, the highest concentration tested for (*R, R*)-GL-586 (20 µM) produced much lower proteome-wide reactivity than the lowest concentration tested for **14** (5 µM) (**Figure 3A**). In the cysteine-directed ABPP experiments, (*R, R*)-GL-373 showed strong engagement (> 75% blockade of iodoacetamide-desthiobiotin (IA-DTB) reactivity) of only three cysteines (NONO_C145; FAM213_C83/85, and USP48_C409), while (*R*)-SKBG-1 engaged > 20 additional cysteines (**Figure 3B** and **Dataset S1**). The lower proteomic reactivity of the CFA-based NONO ligands correlated with decreased glutathione reactivity compared to CA compounds (**Table S2**).

**Figure 3.**
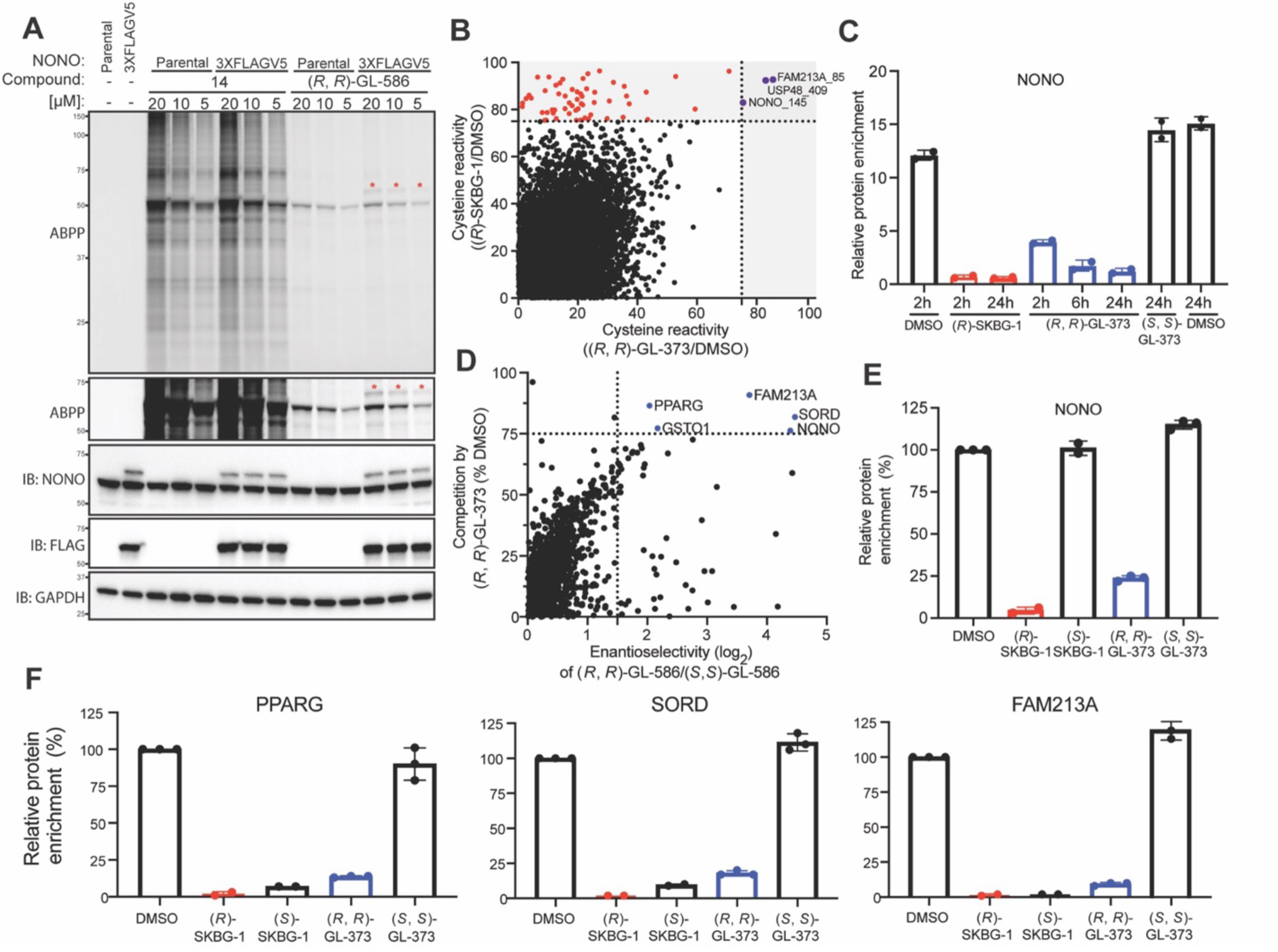
CFA ligands target NONO with improved proteome-wide selectivity. (A) Gel-ABPP data showing concentration-dependent proteome-wide reactivity of alkynes **14** and (*R, R*)-GL-586 (1 h) in parental 22Rv1 cells or 3XFLAGV5-WT-NONO-expressing 22Rv1 cells. Gel-ABPP experiments were performed as described in Figure 2C. Red asterisks (*) highlight signals corresponding to molecular weight of 3XFLAGV5-NONO. Data are from a single experiment representative of two independent experiments. (B) Cysteine-directed ABPP data showing global cysteine reactivity profiles for (*R*)-SKBG-1 and (*R, R*)-GL-373 (20 µM, 6 h) in parental 22Rv1 cells. Data represent a total of 12,017 quantified cysteines, and areas shaded in grey highlight cysteines displaying substantial engagement (>75% blockade of IA-DTB reactivity) by (*R*)-SKBG-1 and/or (*R, R*)-GL-373 (red signals: cysteines engaged by (*R*)-SKBG-1; purple signals: cysteines engaged by both (*R*)-SKBG-1 and (*R, R*)-GL-373). Data are average values from two independent experiments. (C) Protein-directed ABPP data showing time-dependent engagement of NONO by (*R*)-SKBG-1 and (*R, R*)-GL-373 in MCF7 cells. Cells were treated with parent compounds (20 µM) for the indicated times followed by alkyne (*R, R*)-GL-586 (10 µM, 1 h) and analyzed by protein-directed ABPP as described previously.^28^ Data are average values ± SD for two independent experiments. (D) Protein-directed ABPP data comparing the stereoselective enrichment of proteins by (*R, R*)-GL-586 vs (*S, S*)-GL-586 (10 µM, 1 h) from MCF7 cells and the competitive blockade of this enrichment by pre-treatment with (*R, R*)-GL-373 (20 µM, 6 h). Dashed lines mark proteins that showed substantial stereoselective enrichment (log2 > 1.5) by (*R, R*)-GL-586 and substantial blockade of this enrichment (> 75%) by (*R, R*)-GL-373. (E, F) Bar graphs showing protein-directed ABPP data for NONO (E) and additional proteins (F) from (D) that were stereoselectively enriched by (*R, R*)-GL-586 and substantially engaged by (*R, R*)-GL-373. Data are average values ± SD for two-three independent experiments.

We also verified stereoselective engagement of NONO by CFA ligands in a second human cancer cell line – the hormone-sensitive breast cancer line MCF7. In these experiments, we compared the time-dependent engagement of NONO by (*R*)-SKBG-1 and (*R, R*)-GL-373 by protein-directed ABPP^28^ using (*R, R*)-GL-586 as the enrichment probe, which revealed that, while (*R*)-SKBG-1 engaged NONO more rapidly than (*R, R*)-GL-373 (2 h time point, **Figure 3C** and **Dataset S1**), both compounds near-completely engaged NONO at later time points, and this engagement was sustained for at least 24 h (**Figure 3C** and **Dataset S1**), consistent with the long half-life reported previously for NONO (>50 h).^32^ The protein-directed ABPP experiments additionally verified stereoselective engagement of NONO by (*R, R*)-GL-373 (in comparison to (*S, S*)-GL-373; **Figure 3C-E** and **Dataset S1**) along with a handful of additional off-targets (**Figure 3D**). Interestingly, unlike NONO (**Figure 3E**), most of these off-targets were strongly engaged by both (*R*)-SKBG-1 and (*S*)-SKBG-1 (**Figure 3F**), suggesting that the reduced intrinsic electrophilicity of the CFA group, while substantially lowering overall proteomic reactivity, might paradoxically enhance the stereoselective liganding of a subset of proteins that are more indiscriminately engaged by higher reactivity CA compounds.

Our data, taken together, demonstrate that NONO_C145 can be stereoselectively and site-specifically engaged by CFA ligands, and these compounds show much lower proteome-wide reactivity in cells than original CA ligands. We next compared the biological effects of (*R, R*)-GL-373 and (*R*)-SKBG-1 in cancer cells.

### (*R, R*)-GL-373 exhibits NONO-restricted transcriptomic and growth effects in cancer cells

We previously found that (*R*)-SKBG-1 remodeled the transcriptomes of cancer cells, and this effect was largely stereoselective (most of the changing genes were not affected by (*S*)-SKBG-1) and NONO-dependent (most of the changing genes were not affected by (*R*)-SKBG1 in sgNONO cells).^13^ Here, we evaluated the transcriptomic effects of (*R, R*)- and (*S, S*)-GL-373 (20 µM, 6 h) in MCF7 cells by RNA-sequencing (RNA-seq) and found that (*R, R*)-GL-373 produced profound effects on the MCF7 transcriptome that included substantial (log_2_ > 1) decreases in nearly 1000 genes and a more restricted increase in ∼60 genes (**Figure 4A**, **Figure S3**, and **Dataset S2**). These effects were stereoselective (**Figure 4B**, **Figure S4A**, and **Dataset S2**) and generally not observed in sgNONO cells (**Figure 4C**, **Figure S4B**, and **Dataset S2**). Additionally, the NONO-dependent transcriptomic effects of (*R, R*)-GL-373 generally aligned with those of (*R*)-SKBG-1 (5 µM, 6 h) (**Figure 4D** and **Dataset S2**) with the exception of a limited subset of genes that were strongly induced by (*R*)-SKBG-1, but not (*R, R*)-GL-373 (**Figure 4E**). These genes were also induced by i) (*S*)-SKBG-1 and ii) (*R*)-SKBG-1 in sgNONO cells (**Figure 4E**, right panel; and **Figure S4C, D**), suggesting they were NONO-independent effects. We noted that several of the genes induced by (*R*)-SKBG-1, but not (*R, R*)-GL-373, are also regulated by the KEAP1-NRF2 electrophilic/oxidative stress response pathway.^33–35^ We interpret these data to indicate that (*R, R*)-GL-373 causes much less electrophilic stress compared to (*R*)-SKBG-1 and, as a consequence, produces a more NONO-restricted impact on cancer cell transcriptomes.

**Figure 4.**
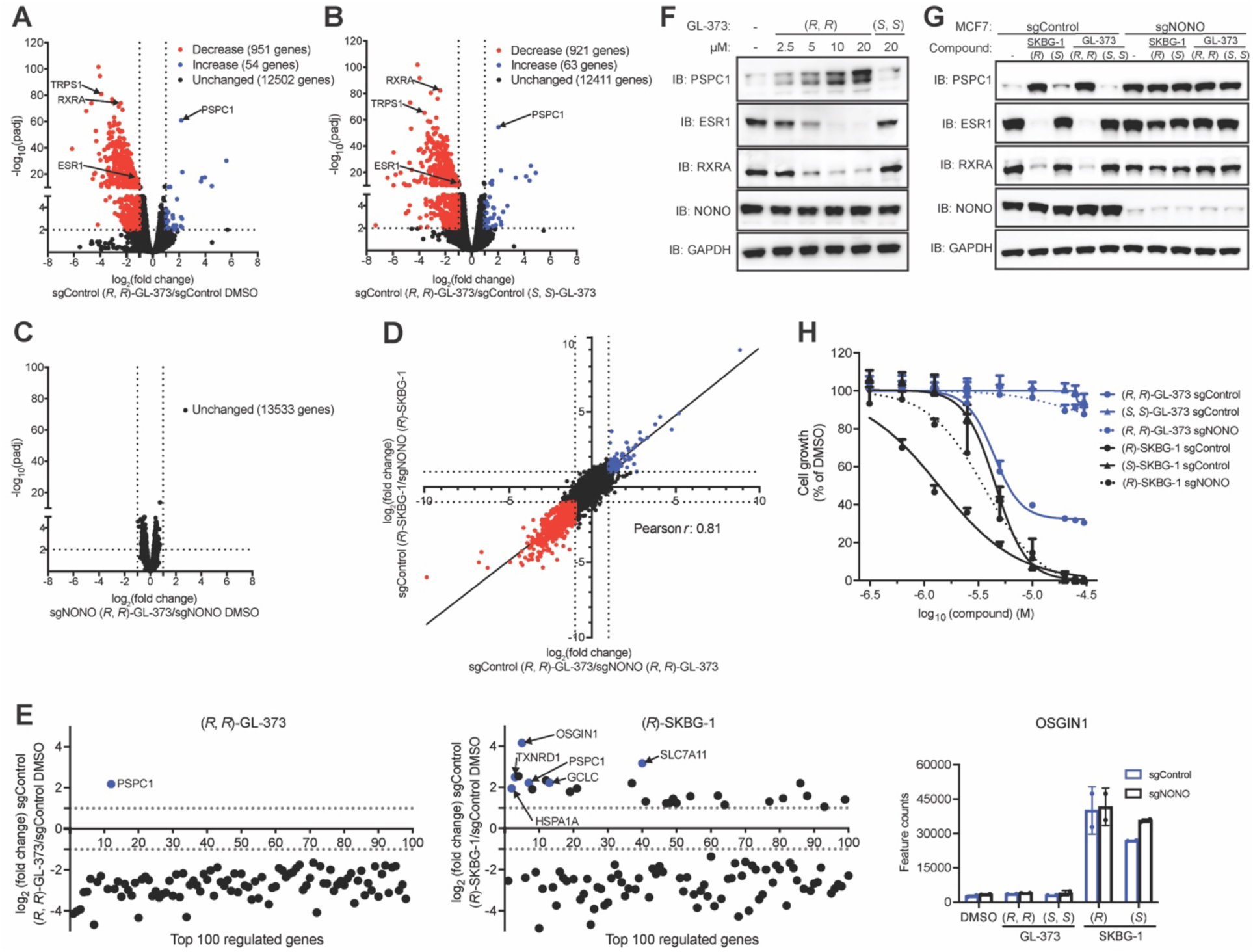
CA ligands produce NONO-restricted transcriptomic and anti-proliferative effects in cancer cells. (A-C) Volcano plots of RNA-seq data showing global gene expression changes (log₂ fold change, L2FC) in MCF7 cells for the following comparison groups: (A) (*R, R*)-GL-373 (20 µM, 6 h) vs DMSO in sgControl cells, (B) (*R, R*)-GL-373 vs (*S, S*)-GL-373 (20 µM each, 6 h) in sgControl cells, and (C) (*R, R*)-GL-373 (20 µM, 6 h) in sgControl vs sgNONO cells. Substantially (|L2FC| > 1) and significantly (padj < 0.01) changing transcripts are highlighted, with red indicating decreased transcripts and blue indicating increased transcripts. (D) Correlation plot comparing the transcriptomic effects of (*R, R*)-GL-373 and (*R*)-SKBG-1 in sgControl vs sgNONO MCF7 cells. (E) Left and middle graphs show RNA-seq changes for the top-100 most significantly altered transcripts in sgControl cells treated with (*R, R*)-GL-373 (20 µM, 6 h) vs DMSO (left graph) or with (*R*)-SKBG-1 (5 µM, 6 h) vs DMSO (middle graph) comparisons. Genes are ordered from left to right based on increasing padj value (x-axis). Highlighted in blue are select transcripts increased by i) both (*R, R*)-GL-373 and (*R*)-SKBG-1 (PSPC1); or ii) only by (*R*)-SKBG-1 that are also regulated by the KEAP-NRF2 electrophilic/oxidative stress response pathway.^33–35^ Right graph shows RNAseq data for OSGIN1, a representative NRF2/KEAP1 pathway-regulated gene that was increased by (*R*)-SKBG-1 and (*S*)-SKBG-1, but not (*R, R*)-GL-373 or (*S, S*)-GL-373, in both sgControl and sgNONO cells. For (A-E), data represent average values of two independent RNA-seq experiments. (F) Western blot analysis showing concentration-dependent effects of (*R, R*)-GL-373 and (*S, S*)-GL-373 on ESR1, RXRA, and PSPC1 protein expression in sgControl MCF7 cells (2.5 – 20 µM compound; 24 h). (G) Western blot analysis comparing effects of the indicated compounds (5 µM for (*R*)-SKBG-1 and (*S*)-SKBG-1; 20 µM for (*R, R*)-GL-373 and (*S, S*)-GL-373; 24 h) on PSPC1, ESR1, and RXRA protein in sgControl vs sgNONO cells. For (F, G), data are from a single experiment representative of at least two independent experiments. (H) Effects of indicated compounds on the growth of sgControl or sgNONO MCF7 cells. Cells were treated with compounds for 48 h, followed by a second treatment at the same concentration, and cell proliferation measured 72 h after the second treatment using the CellTiter-Glo assay. GI_50_ values: (*R*, *R*)-GL-373 in Control: 4.4 µM (95% CI 4.2–4.6 µM; (*S*, *S*)-GL-373 in sgNONO: > 20 µM (95% CI undefined); (*R*, *R*)-GL-373 in sgNONO > 20 µM (95% CI undefined µM); GI_50_ values: (R)-SKBG-1 in sgControl, 1.4 µM (95% CI 1.2–1.6 µM); (*S*)-SKBG-1 in Control, 4.5 µM (95% CI 4.3–4.8 µM); (*R*)-SKBG-1 in sgNONO: 3.7 µM (95% CI 3.1–4.4 µM). Data are average values ± SD for six independent experiments.

The transcriptomic effects of (*R, R*)-GL-373 included the downregulation of several cancer-relevant genes, including those encoding the transcription factors ESR1, RXRA, and TRPS1 (**Figure 4A-C**). (*R, R*)-GL-373 also increased the expression of the NONO paralog PSPC1 (**Figure 4A, B, E**), an effect that is also observed in cells genetically disrupted for NONO^13,36^ (and has been proposed to reflect a compensatory response to loss of NONO.^15,36^ Western blotting experiments performed at 24 h post-treatment confirmed that (*R, R*)-GL-373 additionally induced stereoselective and NONO-dependent decreases in the abundance of ESR1 and RXRA proteins, as well as increases in PSPC1 (**Figure 4F, G**). These effects were observed across a concentration range of 5-20 µM of (*R, R*)-GL-373 with near complete loss of ESR1 in MCF7 cells treated with 10 µM (**Figure 4F**).

Finally, both (*R*)-SKBG-1 and (*R, R*)-GL-373 produced stereoselective and NONO-dependent reductions in the proliferation of MCF7 cells (**Figure 4H**). While (*R*)-SKBG-1 showed higher potency than (*R, R*)-GL-373 in these cell growth assays, the antiproliferative effects of (*R, R*)-GL-373 showed much greater stereoselectivity and NONO-dependency, likely reflecting the lower general electrophilic stress caused by this compound compared to (*R*)-SKBG-1.

These data, taken together, indicate that CFA ligands exhibit NONO-dependent effects on the transcriptomes and growth of cancer cells while avoiding the NONO-independent electrophilic stress caused by original CA ligands.

## Discussion

RNA-binding proteins (RBPs) represent an exciting opportunity for chemical biology and drug discovery.^11,12^ A substantial portion of the human proteome binds RNA as a primary or secondary function,^2,37^ and many RBPs have clear human disease-relevance.^38^ Additionally, drugs such as risdiplam for the treatment of spinal muscular atrophy have shown the potential for RBPs to serve as targets that promote therapeutic gene expression.^8^ Nonetheless, most RBPs lack chemical probes, likely reflecting, at least in part, the technical challenges associated with screening this class of proteins, which are often parts of large and dynamic complexes. ABPP and related chemical proteomic methods offer an attractive way to identify ligands for RBPs directly in native cellular environments, as we recently demonstrated with the discovery of CAs that covalently bind NONO to remodel the transcriptomes of cancer cells.^13^ However, the structural basis for CA binding to NONO, as well as the potential to advance this interaction into more selective chemical probes, have remained important open questions.

The (*R*)-SKBG-1-NONO structure provides clear evidence of a small molecule-binding pocket proximal to the RRM1 and RRM2 RNA-binding domains. Our biochemical data measuring NONO binding affinity for RNAs also support previous cell-based (eCLIP) experiments^13^ that covalent binding of (*R*)-SKBG-1 to this pocket enhances NONO-RNA interactions. How precisely this outcome is achieved remains unclear. Limited structural information is available on DBHS protein-RNA complexes, and the only available structure of RNA-bound SFPQ homodimers^22^ would suggest that (*R*)-SKBG-1 might disrupt NONO-RNA interactions. On the other hand, the various structures of DBHS proteins show highest variability in the pose of the RRM1 domain,^15^ and it is possible that (*R*)-SKBG-1 reactivity with C145 strengthens RNA binding to NONO by resembling a productive conformation for RNA interactions. A structure of a ternary covalent ligand:NONO:RNA complex may be required to more fully understand how small molecule reactivity with C145 stabilizes NONO-RNA interactions.

The (*R*)-SKBG-1-NONO structure also revealed a well-aligned binding pose for C145 reactivity with the CA α-carbon of (*R*)-SKBG-1, which inspired the generation of a less electrophilic CFA analog (*R, R*)-GL-373. While (*R, R*)-GL-373 showed some loss (∼four-fold) in potency for engaging NONO, the much greater proteome-wide selectivity of this interaction resulted in profound NONO-dependent remodeling of breast cancer cell transcriptomes without the induction of general electrophile stress caused by (*R*)-SKBG-1. These data thus provide further evidence for the utility of the CFA as a tempered reactive group in the development of covalent chemical probes.^25–27,39^ We should acknowledge, however, that the cellular activity of (*R, R*)-GL-373 likely benefited from the long half-life of NONO, which allowed for near-complete and sustained engagement of C145 over a 24 h time period (**Figure 3C**). Shorter half-life proteins may require higher potency CFA compounds to achieve similar levels of engagement in cells. (*R, R*)-GL-373 also exhibited stereoselective reactivity with a handful of proteins beyond NONO, which reinforces the importance of performing additional control experiments in NONO-disrupted (sgNONO) cells to verify on-target (NONO-dependent) activities, as we demonstrated herein for the transcriptomic and cell growth effects of (*R, R*)-GL-373.

Looking forward, we are intrigued by the impact of covalent NONO ligands on the expression of breast cancer-relevant transcription factors like ESR1 and TRPS1, which mirror the previously reported effects of NONO ligands on androgen receptor expression in prostate cancer cells.^13^ From a translational perspective, these findings suggest that targeting NONO may offer a way to suppress lineage drivers in breast and prostate cancer that is complementary to the targeted protein degradation strategies under investigation in the clinic.^40,41^ However, the current NONO ligands, including (*R, R*)-GL-373, suppress a large swath of transcripts, which results in a more general antiproliferative effect.^13^ Using the (*R*)-SKBG-1-NONO co-crystal structure as a guide, we speculate that it may be possible to develop ligands with improved transcript selectivity by, for instance, modifying the structures of these compounds to interface more directly with RNA. Finally, we also anticipate that the tempered reactivity and greater proteomic specificity displayed by (*R, R*)-GL-373 will facilitate addressing fundamental mechanistic questions that include: 1) how do NONO ligands promote such a rapid loss of transcripts in cancer cells? 2) what features do these transcripts share in common that confer sensitivity to NONO ligands? 3) how do cells exposed to NONO ligands upregulate paralogous DBHS proteins?; and 4) what cellular factors confer sensitivity or resistance to NONO ligands? Considering the diverse transcriptional and post-transcriptional roles performed by DBHS proteins in human cells, we believe the development of (*R, R*)-GL-373 will serve as a useful tool for studying the functions of this important family of RBPs.

### Limitations of the study

Given that our (*R*)-SKBG1:NONO co-crystal structure lacks RNA, we can only speculate, at this stage, on how covalent ligands strengthen NONO interactions with RNA. Future efforts to determine a ternary ligand:NONO:RNA structure hold promise to provide more detailed mechanistic insights. Despite exhibiting substantial improvements in proteome-wide selectivity, CFA ligands like (*R, R*)-GL-373 still show only low-µM engagement of NONO. While this degree of potency does not hinder the use of (*R, R*)-GL-373 in cell biological studies, improvements will be needed before such CFA ligands can be applied to study NONO function in vivo. From a translational perspective, covalent NONO ligands suppress the expression of key cancer drug targets like ESR1, but this effect is also accompanied by decreases in a large array of other transcripts that will likely limit the cell type-specificity of the antiproliferative activity of NONO ligands. Whether NONO ligands can be optimized to show transcript-restricted activity remains an important unanswered question.

### Significance

RNA-binding proteins (RBPs) are a large and diverse class of proteins that contribute to virtually all steps of the transcriptional and post-transcriptional regulation of gene expression. Chemical tools are lacking for RBPs, which often form dynamic complexes in cells that are challenging to reconstitute for conventional small-molecule screening. Activity-based protein profiling (ABPP) and related chemical proteomic methods offer a potentially powerful way to discover small molecule ligands for RBPs, as we have shown in the identification of covalent ligands targeting the RBP NONO. Here, we describe the structural characterization of a ligandable pocket in proximity to the RNA-binding domain of NONO that offers mechanistic hypotheses for covalent compound-induced stabilization of NONO-RNA interactions. In developing chlorofluoroacetamide (CFA) ligands for NONO, we further show the potential for targeting this RBP with highly selective covalent probes that lead to the global remodeling of cancer cell transcriptomes and suppression of cancer cell growth without causing general oxidative stress associated with higher reactivity electrophilic compounds. In summary, our findings provide a powerful new chemical tool to study NONO-mediated regulation of the transcriptome and, through doing so, highlight the broader potential for chemical proteomics to address the ligandability of RBPs.

## Resource Availability

**Lead contact.** Further information and requests for resources and reagents should be directed and will be fulfilled by lead contact, Benjamin F. Cravatt.

## Materials availability

All compounds generated in this study are available from the lead contact with a completed materials transfer agreement.

## Data and code availability

- The mass spectrometry proteomics data have been deposited to the ProteomeXchange Consortium via the PRIDE95 partner repository with the dataset identifier PXD064685. Processed proteomic data are provided in **Dataset S1**. The RNAseq data have been deposited at the Gene Expression Omnibus (GEO) at the NCBI with the identifier GSE299099. All other data that support the findings of this study and codes are available from the corresponding authors upon reasonable request. All data are publicly available as of the date of publication.
- This paper does not report original code.
- Any additional information required to reanalyze the data reported in this paper is available from the lead contact upon request.

## Supporting information

PDB Validation Report

## Acknowledgements

This work was supported by the NIH (R35 CA231991; R01 CA238249) and delivered as part of the eDyNAmiC team supported by the Cancer Grand Challenges partnership funded by Cancer Research UK (CGCATF-2021/100012 + CGCATF-2021/100021) and the National Cancer Institute (OT2CA278688 + OT2CA278692). This research was undertaken in part using the MX2 beamline at the Australian Synchrotron, part of ANSTO, and made use of the Australian Cancer Research Foundation (ACRF) detector. Australian Research Council [DP220103667 to C.S.B. and A.H.F.; LE120100092, LE140100096, LE230100156 to C.S.B.; DE240101210 to A.C.M.; FT180100204 to A.H.F.], National Health and Medical Research Council of Australia [APP1147496 to C.S.B. and A.H.F.] and the Cancer Research Trust [Australian Centre for RNA Therapeutics in Cancer, A.H.F. and C.S.B.].

## Author contributions

G.L.L., B.F.C., and C.S.B. conceived the study. G.L.L generated the proteomic data. G.L.L, B.M., and B.F.C. performed analysis of the proteomic data. G.L.L. performed gel-based profiling. T.K.H. crystallized and solved the structure of NONO:(R)-SKBG-1. A.V.G. designed and performed the microscale thermophoresis assay. G.L.L performed cell growth assay. G.L.L prepared RNA-seq samples. G.L.L. and C.T.F performed western blotting. G.L.L, B.M., W.H.B., and B.F.C., performed processing of sequencing data and/or analysis of RNA-seq data. B.M. supervised compound characterization. A.C.M., J.G, G.W.Y., and A.H.F provided support and resources. G.L.L., B.M., B.F.C., and C.S.B. wrote and edited the manuscript. B.F.C. and C.S.B. supervised this study.

## Supplementary information

Detailed methods are provided in the online version of the paper and include the following:

- **KEY RESOURCE TABLE**
- **EXPERIMENTAL MODEL AND STUDY PARTICIPANT DETAILS**

- Cell lines
- **METHOD DETAILS**

- Crystallography
- Microscale thermophoresis assay
- Plasmid amplification and purification
- Mutagenesis
- Plasmids for stable expression
- NONO knockout by CRISPR/Cas9
- Generation of 22Rv1 NONO C-FLAG WT or C145S cell line
- Western blot analysis
- Gel-based ABPP
- Cysteine-directed ABPP
- HPLC fractionation
- Protein-directed ABPP
- High-pH spin column fractionation
- TMT LC-MS Analysis
- MS data processing
- Data analysis – Cysteine-directed ABPP
- Data analysis – Protein-directed ABPP
- RNA-seq sample preparation
- RNA-seq library preparation with polyA selection and HiSeq Sequencing
- RNA-seq data processing and quantification
- RNA-seq data analysis
- Cell growth inhibition assay
- Chemistry

---

**Figure S1.**
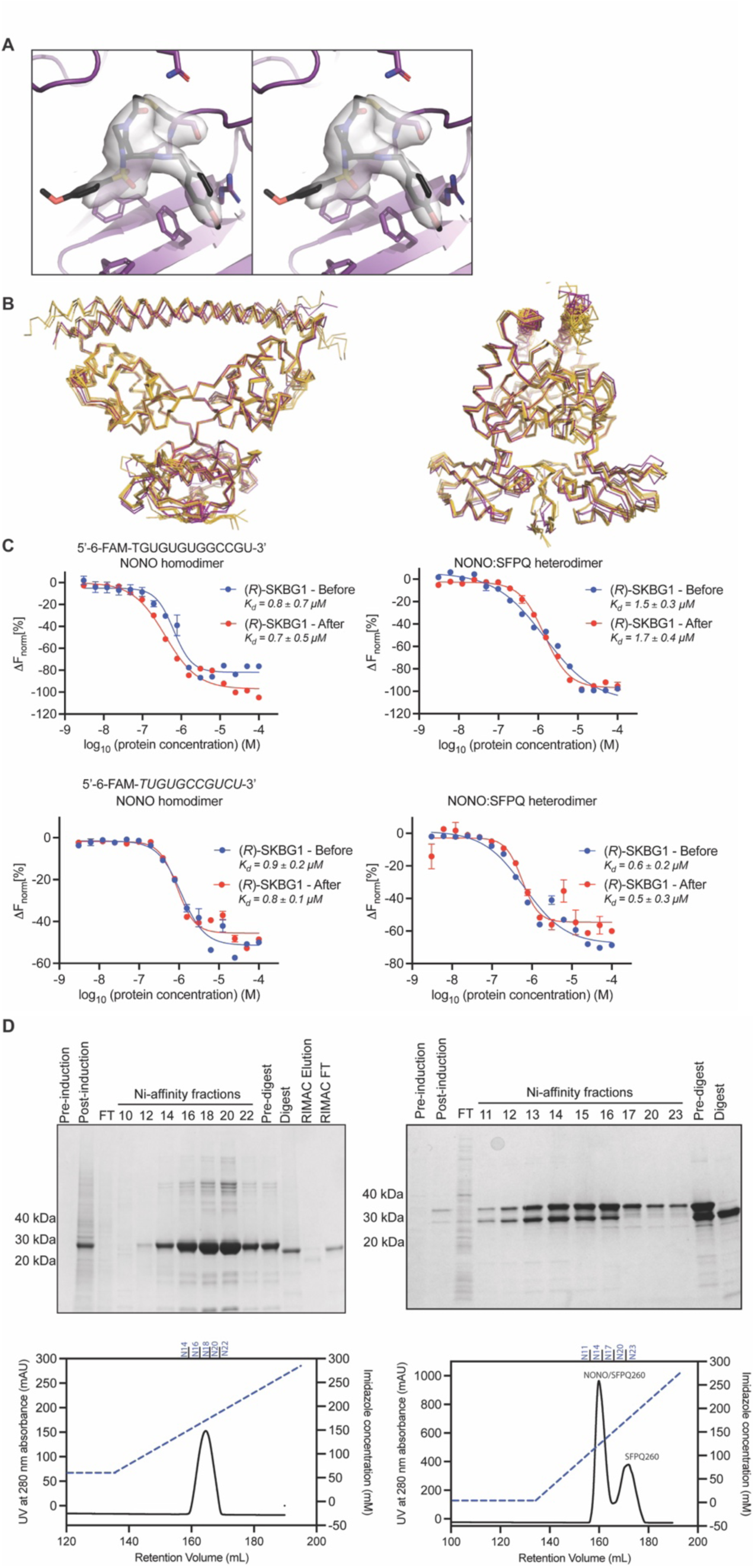
Structural characterization of (*R*)-SKBG-1-bound-NONO homodimer (related to. Figure 1). (A) Stereoview (wall-eyed) of chain A of NONO (purple cartoon and sticks) adducted by (*R*)-SKBG-1 (black sticks), superimposed on an *m*Fo-*D*Fc, α_calc_ electron density map (3σ, grey surface) calculated omitting C145 and (*R*)-SKBG-1 from chain A. The two images shown are stereopairs. (B) Superposition of the two (*R*)-SKBG-1:NONO homodimers in this work (PDB 9NZI; purple) with the six apo NONO homodimers observed in PDB 5IFM (yellow), reveals structural similarity (RMSD < 1 Angstrom), with some diversity in the NOPS and coiled coil regions. (C), Microscale thermophoresis data for two FAM-labeled RNA oligonucleotides interacting with either a NONO homodimer or NONO-SFPQ heterodimer showing that the order of addition of (*R*)-SKBG-1 (before or after RNA) to purified protein did not impact affinity measurements. Data are average values ± SD for three independent experiments. (D) Nickel affinity chromatography with SDS-PAGE demonstrates production of NONO homodimer when only NONO is expressed, and separable samples of NONO/SFPQ heterodimer and SFPQ homodimer when NONO and SFPQ are coexpressed. FT indicates flow-through sample. Fractions collected are labeled on both chromatogram and gel.

**Figure S2.**
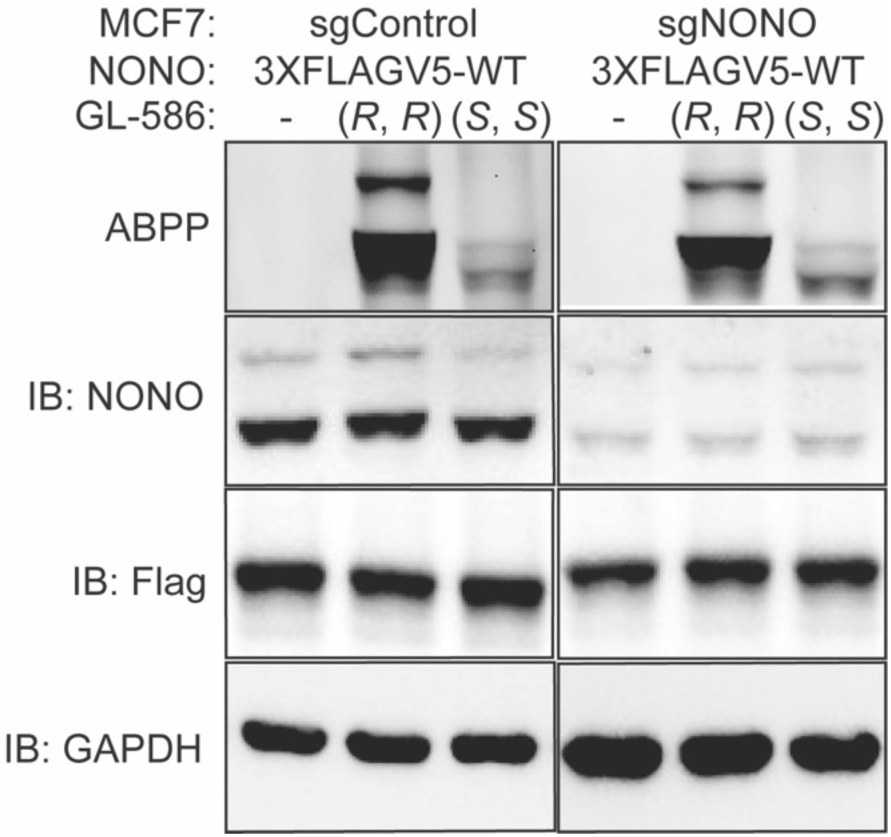
Chlorofluoroacetamide (CFA) ligands stereoselectively and site-specifically engage NONO (related to. Figure 2). Gel-ABPP data showing the proteomic reactivity of alkynes (*R, R*)-GL-586 and (*S, S*)-GL-586 (10 *µ*M, 1 h) in 22Rv1 cells stably expressing 3X-FLAG-V5 epitope-tagged WT-NONO [3XFLAGV5]. 3XFLAGV5 (sgControl) cells also contain endogenous NONO, whereas 3XFLAGV5 (sgNONO) cells have NONO genetically deleted by CRISPR/Cas9 (see NONO immunoblot for degree of NONO disruption in the sgNONO cells). GL-586-reactive proteins were detected by CuAAC conjugation of an azide-rhodamine reporter group, followed by SDS–PAGE, and in-gel fluorescence scanning.

**Figure S3.**
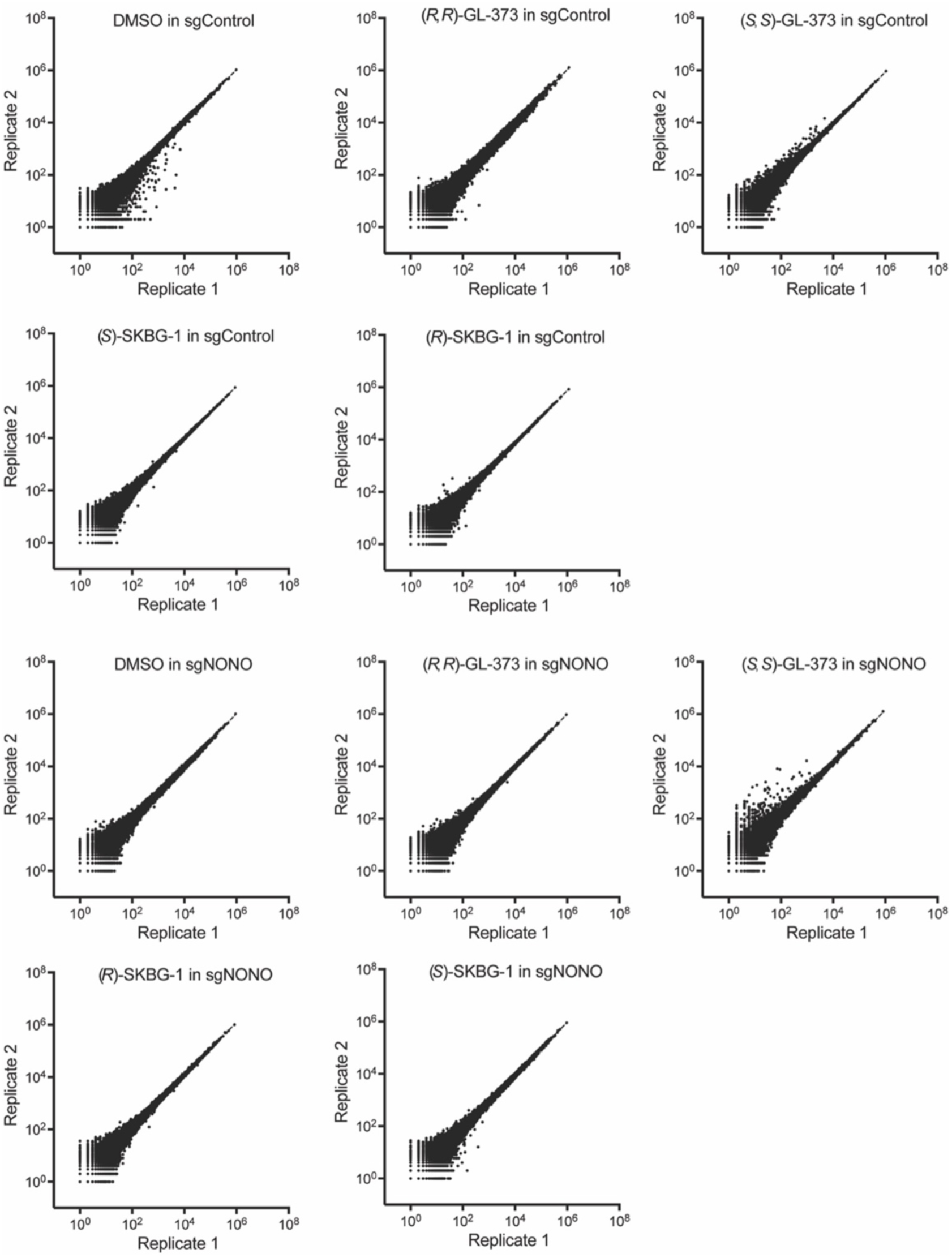
Correlation plots for replicate RNA-seq experiments (related to Figure 4). Plots depicting replicate feature count agreement for each treatment condition: DMSO, (*R*)-SKBG-1 (5 µM), and (*S*)-SKBG-1 (5 µM), (*R, R*)-GL-373 (20 µM), (S, S)-GL-373 (20 µM), in both sgControl and sgNONO MCF7 cells treated for 6 h.

**Figure S4.**
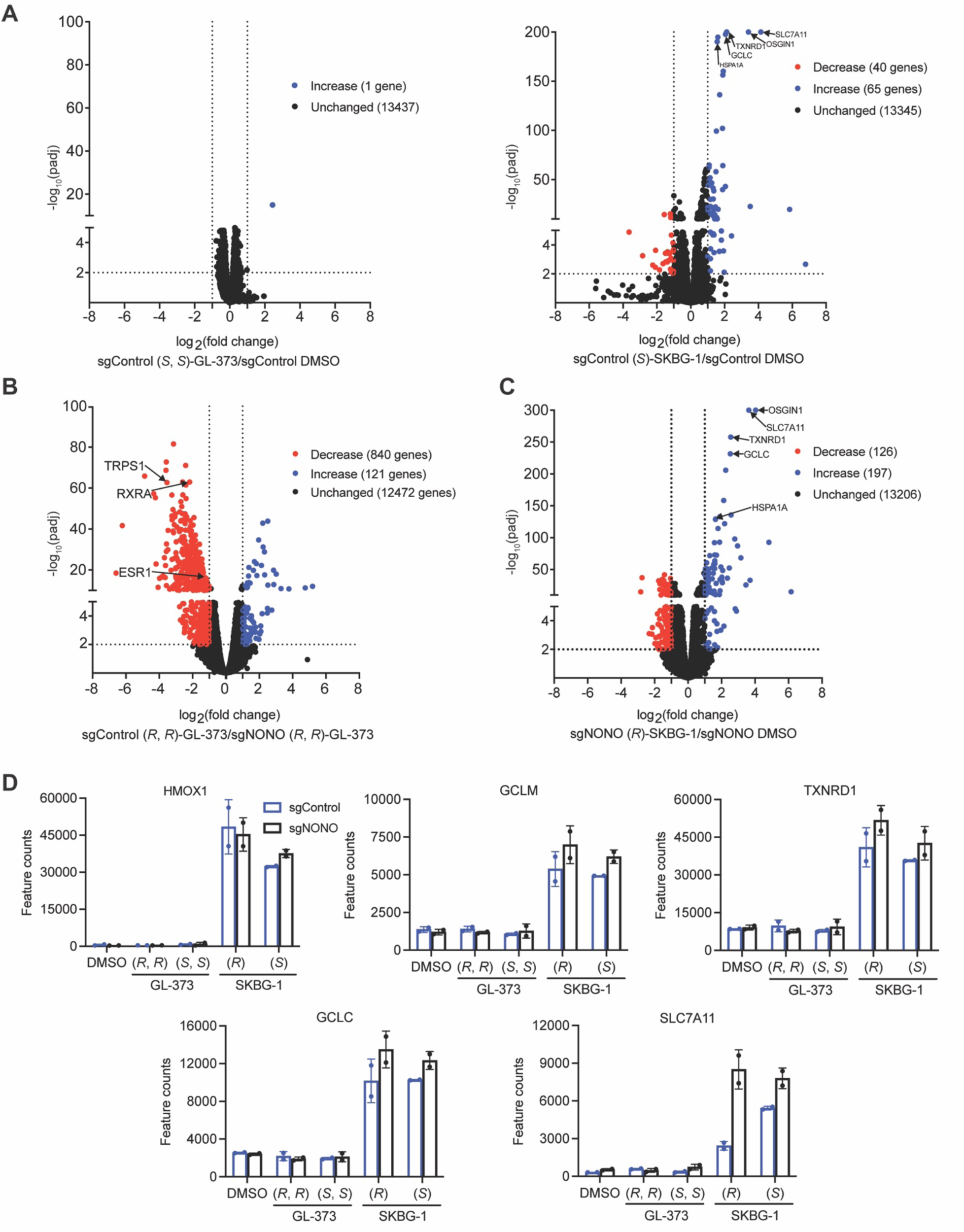
CA ligands produce NONO-restricted transcriptomic and anti-proliferative effects in cancer cells (related to. Figure 4). (A) Volcano plots of RNA-seq data showing global gene expression changes (log₂ fold change, L2FC) in MCF7 cells for the following comparison groups: Left, (*S, S*)-GL-373 (20 µM, 6 h) vs DMSO in sgControl cells; right, (*S*)-SKBG-1 (5 µM, 6 h) vs DMSO in sgControl cells. Substantially (|L2FC| > 1) and significantly (padj < 0.01) changing transcripts are highlighted, with red indicating decreased transcripts and blue indicating increased transcripts. (B, C) Volcano plots of RNA-seq data showing global gene expression changes (log₂ fold change, L2FC) in MCF7 cells for the following comparison groups: (B), (*R, R*)-GL-373 (20 µM, 6 h) in sgControl cells vs *R, R*)-GL-373 (20 µM, 6 h) in sgNONO cells; (C) (*R*)-SKBG-1 (5 µM, 6 h) vs DMSO in sgNONO cells. Substantially (|L2FC| > 1) and significantly (padj < 0.01) changing transcripts are highlighted, with red indicating decreased transcripts and blue indicating increased transcripts. (D) Bar graphs illustrating expression levels of additional genes regulated by the KEAP-NRF2 electrophilic/oxidative stress response pathway across all treatment conditions. Data are average values ± SD for two independent experiments.

**Table S1.** Data collection and refinement statistics for (*R*)-SKBG1-NONO homodimer co-crystal structure. ^☨^Calculated using MolProbity.

| Data & Refinement Statistics |  |
| --- | --- |
| Wavelength (Å) | 0.9537 |
| Space group | P 1 2<sub>1</sub> 2 |
| Unit-cell parameters | a = 61.1 Å, b = 83.0 Å, c = 117.2 Å |
| | $\alpha = 94.5^\circ$, $\beta = 89.9^\circ$, $\gamma = 103.6^\circ$ |
| Resolution Range (Å) | 48.31 – 2.52 |
| $R_{merge}$ | 0.063 (1.205) |
| $\langle I/\sigma(I) \rangle$ | 11.6 (1.2) |
| $CC_{1/2}$ | 0.998 (0.479) |
| Completeness (%) | 99.3 (97.2) |
| Refinement |  |
| Unique reflections | 40 730 (3954) |
| $R_{work}$ | 0.225 |
| $R_{free}$ | 0.268 |
| B-factors (Å<sup>2</sup>) |  |
| Protein | 92.3 |
| Ligand | 120.1 |
| Water | 115.8 |
| Ramachandran Plot† |  |
| Favoured region (%) | 99.4 |
| Disallowed region (%) | 0.63 |
| R.m.s. deviations |  |
| Bond length (Å) | 0.0016 |
| Bond angles (°) | 0.40 |

**Table S2.** Glutathione reactivity values for CA and CFA compounds. Summary of the rate (M^-1^•s^-1^) glutathione consumption by representative stereoprobes at 2 h.

| Compound | $M^{-1}\cdot s^{-1}$ | Timepoint (min) |
| --- | --- | --- |
| (R)-SKBG-1 | 0.187 | 120 |
| (S)-SKBG-1 | 0.177 | 120 |
| (S, R)-GL-373 | 0.058 | 120 |
| (R, R)-GL-373 | 0.061 | 120 |
| (S, S)-GL-373 | 0.054 | 120 |
| (R, S)-GL-373 | 0.055 | 120 |

