## Supplementary Information for "Structural and mechanistic analysis of covalent ligands targeting the RNA-binding protein NONO"

### Reagents and materials

| Reagent/Material | Source | Identifier |
|---|---|---|
| Antibodies | | |
| Rabbit anti- ESR1 Receptor | Cell Signaling Tech | Cat#: 5153S |
| RXRA Rabbit Poly Ab | Proteintech | Cat#: 21218-1-AP |
| Mouse anti- NONO | BD Biosciences | Cat#: 611278 |
| Rabbit anti- NONO | Bethyl Laboratories | Cat#: A300-587A |
| Anti-GAPDH HRP | Santa Cruz Biotechnology | Cat#: sc-47778 HRP |
| Mouse anti- PSPC1 | Sigma-Aldrich | Cat#: SAB4200503 |
| Mouse anti- FLAG | Sigma-Aldrich | Cat#: F1804 |
| HRP-labeled anti-mouse | Cell Signaling Tech | Cat#: 7076S |
| HRP-labeled anti-rabbit | Cell Signaling Tech | Cat#: 7074S |
| Cell culture | | |
| RPMI-1640 media | Corning | Cat#: 15-040-CV |
| DMEM media | Corning | Cat#: 15-013-CV |
| Glutamax | Life Technologies | Cat#: 35050061 |
| Penicillin-Streptomycin | Lonza | Cat#: 17-603E |
| Fetal bovine serum | Omega Scientific | Cat#: FB-21 |
| PEI MAX Linear MW 40,000 | Polysciences | Cat#: 24765-1 |
| RNAiMax | ThermoFisher | Cat#: 13778030 |
| Fugene 6 | Promega | Cat#: E2691 |
| Polybrene | Santa Cruz | Cat#: 134220 |
| Blasticidin | Fisher Scientific | Cat#: 50712728 |
| Puromycin | Sigma-Aldrich | Cat#: P8833 |
| Bacterial strains | | |

| | | |
|---|---|---|
| One Shot Stbl3 Chemically Competent E. coli | Thermo Scientific | Cat#: C737303 |
| ccdB Survival T1 | Invitrogen | Cat#: 11828029 |
| Chemicals and materials | | |
| Pierce ECL Western Blotting Substrate | ThermoFisher | Cat#: 32106 |
| SuperSignal West Femto PLUS Chemiluminescent Substrate | ThermoFisher | Cat#: PI34095 |
| Novex 10% Tris-Glycine Mini Gels | Life Technologies | Cat#: XP00105BOX |
| Nitrocellulose western blotting membrane, 0.45 mM | GE Healthcare Amersham | Cat#: 10600002 |
| Complete+Ultra Mini EDTA-Free Protease Inhibitor Cocktail Tablets | Roche | Cat#: 05892791001 |
| DMSO | Corning | Cat#: 25-950-CQC |
| Carbenicillin | Fisher | Cat#: NC0753434 |
| Chloramphenicol | Sigma-Aldrich | Cat#: C0378 |
| Spectinomycin | Sigma-Aldrich | Cat#: S4014 |
| Desthiobiotin polyethyleneoxide iodoacetamide | Santa Cruz Biotechnology | Cat#: sc-300424 |
| Urea | Fisher Scientific | Cat#: M1084871000 |
| Iodoacetamide | Sigma-Aldrich | Cat#: I1149-25G |
| Dithiothreitol (DTT) | Fisher Bioreagents | Cat#: BP172-25 |
| Tris(benzyltriazolylmethyl)amine (TBTA) | TCI | Cat#: T2993 |
| Copper(II) sulfate, anhydrous | Sigma-Aldrich | Cat#: 451657-10G |
| Tris(2-carboxyethyl)phosphine HCl (TCEP) | Sigma-Aldrich | Cat#: 75259 |
| Biotin-PEG4-azide | Chempep | Cat#: 271606 |
| Sequencing grade modified trypsin | Promega | Cat#: V5111 |
| Lys-C, Mass Spec Grade | Promega | Cat#: VA1170 |

| | | |
|---|---|---|
| Streptavidin agarose resin | Fisher Scientific | Cat#: 20353 |
| Tween-20 | Fisher Bioreagents | Cat#: BP337-500 |
| Triton X-100 | EMD Millipore | Cat#: TX1568 |
| Nonidet P40 substitute (Igepal CA-630) | USB Corporation | Cat#: 19628 |
| EPPS | Sigma-Aldrich | Cat#: E0276 |
| Calcium carbonate | Fisher | Cat#: C77 |
| SDS | Fisher | Cat#: BP166 |
| $K_2CO_3$ | Fisher | Cat#: P208 |
| Triethylammonium bicarbonate buffer | Sigma-Aldrich | Cat#: T7408-500ML |
| TMT10plex$^{TM}$ | ThermoFisher | Cat#: 90406 |
| TMTpro$^{TM}$ 16plex tag | ThermoFisher | Cat#: A44520 |
| Acetonitrile | VWR Intl | Cat#: BJAS017-0100 |
| Hydroxylamine solution | Sigma-Aldrich | Cat#: 467804-10ML |
| Formic acid, ~98%, for mass spectrometry | Honeywell Fluka | Cat#: 94318-250ML-F |
| Bovine serum albumin | Sigma-Aldrich | Cat#: A2153 |
| Sep-Pak C18 cartridges | Waters | Cat#: WAT054955 |
| Spin Columns, Desalting, Pierce Peptide | ThermoFisher | Cat#: PI89852 |
| Power Blotter Select Transfer Stacks. Nitrocellulose | Thermo Scientific | Cat#: PB3210 |
| SuperSignal West Pico PLUS Chemiluminescent Substrate T | Thermo Scientific | Cat#: 34580 |
| Extraction disks, C18 sorbent (for Stage-tip) | 3M empore | Cat#:143863 |
| Commercial assays | | |
| Micro BCA Protein Assay Kit | Thermo Scientific | Cat#: 23235 |
| Cell Titer Glo | Promega | Cat#: 134220 |
| Deposited data | | |

| | | |
|---|---|---|
| Proteomics | This paper | PRIDE: PXD064685 |
| RNA-Sequencing | This paper | GEO: GSE299099 |
| Oligonucleotides | | |
| Primers used in this study | IDT | Methods below |
| Software and algorithms | | |
| MO.Affinity Analysis Software | Nanotemper | https://shop.nanotempertech.com/software/analysis-software/ |
| MOLREP | Version 11.0 /22.07.2010/ | https://www.ccp4.ac.uk/html/molrep.html |
| CCP4 Suite | Version 9 | https://www.ccp4.ac.uk/download/#os=macos |
| COOT | GTK4: 1.1.14 | https://www2.mrc-lmb.cam.ac.uk/personal/pemsley/coot/ |
| PHENIX | Version 1.21 | https://phenix-online.org/ |
| RAW Converter | Version 1.1.0.22; 2004 release | http://fields.scripps.edu/rawconv/ |
| Integrated Proteomics Pipeline (IP2) and ProLuCID | Integrated Proteomics Applications | http://goldfish.scripps.edu/ |
| Prism (v10.4.1) | GraphPad Software | http://www.graphpad.com/scientific-software/prism/ |
| STAR aligner (v2.7.9a) | Cold Spring Harbor Laboratory | https://github.com/alexdobin/STAR |
| R (v2025.05.0+496) | R Core Team | https://www.r-project.org/ |
| Salmon (v1.3.0) | Stony Brook University | https://github.com/COMBINE-lab/salmon |
| DESeq2 (v1.30.1) | European Molecular Biology Laboratory | https://bioconductor.org/packages/release/bioc/html/DESeq2.html |
| Other kits | | |
| Quikchange | Agilent | Cat#: 200521 |
| Gateway Vector Conversion System | Invitrogen | Cat#: 11828029 |
| NEBNext Ultra RNA Library Prep | NEB | Cat#: E7770S |

| | | |
|---|---|---|
| RNeasy Plus kit | QIAGEN | Cat#: 74034 |
| QIAshredder columns | QIAGEN | Cat#: 79654 |
| QIAprep Spin Miniprep kit | QIAGEN | Cat#: 27104 |
| PierceTM BCA Protein Assay Kit | ThermoFisher | Cat#: 23225 |

### Resource Availability

#### Lead Contact

Further information and requests for resources and reagents should be directed to and will be fulfilled by the Lead Contacts, Benjamin F. Cravatt and Charles S. Bond.

#### Materials Availability

All chemical probes and other elaborated electrophilic compounds generated in this study are available from the Lead Contact with a completed Materials Transfer Agreement.

#### Data and Code Availability

Raw proteomic data been deposited to the ProteomeXchange Consortium via the PRIDE partner repository with the dataset identifier PXD064685. Raw RNA-sequencing data has been deposited to NCBI under GEO: GSE299099. Processed proteomic data is provided in Supplementary Datasets 1, RNA-sequencing data is provided in Supplementary Dataset 2.

### Experimental Details

#### Crystallography

Truncated versions of NONO (residues 53-312) homodimer, SFPQ (276-535) homodimer and NONO/SFPQ heterodimer were expressed, purified and concentrated for crystallography and microscale thermophoresis (MST) experiments as described.[1-3]

For crystallization, NONO homodimer (3.7 mg/mL) was incubated with 4x molar excess of (*R*)-SKBG-1 (100 μM in DMSO) for 15 minutes at room temperature. Sitting drop vapour diffusion screens (ProPlex Screen MD1-38, SG1 Screen MD1-88, Pact Premier, JGSG Plus (Molecular Dimensions), Natrix 1 & 2 (Hampton Research) were set up in Intelli-Plate 96-3 (Art Robbins) plates using an Art Robbins Phenix liquid handling robot (reservoir 80 μl, protein:reservoir ratios 1:1, 2:1, 1:2), incubated at 20°C and observed using Formulatrix RockImager. Diffraction quality crystals were obtained from conditions optimized to 0.1 M HEPES, 28% PEG 2000 monomethyl ether. Crystals were cryoprotected by addition of 10% glycerol, harvested and cryopreserved for data collection at the MX2 beamline of the Australian Synchrotron.[4] Complete data to 2.5 Å, reduced with XDS[5] yielded a dataset suitable for structure solution and refinement (details in Supp Table 1). The structure was solved by molecular replacement (MOLREP)[6], CCP4 Suite[7] using a NONO homodimer model derived from chains A and B of PDB entry 5IFM.[8] Subsequent rounds of model fitting, refinement and validation in COOT[9] and PHENIX,[10] including geometric parameters for the C145-linked SKBG-1 (ELBOW),[11] yielded a final model with two NONO homodimers in the asymmetric unit and an R-factor of 22.5% and R-free of 26.2%. The validated structure was deposited in the PDB with code 9NZI.

**Microscale thermophoresis assay (MST)**

For MST experiments, RNA oligonucleotides (5'–6 FAM–TGUGUGUGGCCGU–3' and 5'–6 FAM–TUGUGCCGUCU–3') were purchased from IDT. NONO homodimer, SFPQ homodimer and NONO/SFPQ heterodimer were incubated with a 4x excess (per protein monomer) of (R)-SKBG-1 (100 µM in DMSO), (S)-SKBG-1 or DMSO only. MST was carried out in triplicate on a dilution series of protein in the presence of 200 nM RNA using a Monolith NT.155 instrument, and data analysed using MO. Affinity (NanoTemper). To test the significance of order of addition of SKBG-1 and RNA, measurements were carried out on series of simples of identical composition, but with RNA added first.

**Cell lines**

22Rv1 (CVCL_1045, Male, Prostate carcinoma) and MCF7 (CVCL_0031, Female, Breast adenocarcinoma) cells were obtained from ATCC and grown and maintained in RPMI 1640 media (Thermo Fisher Scientific) containing 10% (v/v) FBS (Omega Scientific), 100 U/mL penicillin-streptomycin (GE Life Sciences), and Glutamax [22Rv1] or DMEM (Thermo Fisher Scientific) containing 10% (v/v) FBS (Omega Scientific), Glutamax, and 100 U/mL penicillin-streptomycin (GE Life Sciences) [MCF7] at 37 °C with 5% $CO_2$. All cell lines were routinely inspected for mycoplasma contamination.

**Cloning and mutagenesis**

**Plasmid amplification and purification**

All plasmids were amplified in *Stbl3* cells (ThermoFisher) using vendor's procedures and purified by miniprep (QIAGEN), except for the pRK5 and pLenti6.2-ccdB-3xFLAG-V5 Gateway destination vectors, which were amplified in ccdB Survival T1 cells (Invitrogen).

**Mutagenesis**

Site-directed mutagenesis was performed on the pDONR 223-NONO construct from the Human ORFome V8.1 Library (Dharmacon) using the Quikchange mutagenesis kit (Agilent). All mutations were verified by DNA sequencing. Primers for mutagenesis (mutated nucleotide in bold and underlined):

NONO-C145S-fwd (T→A): GTGTGCGCTTTGCCAGCCATAGTGCATCC

NONO-C145S-rev: GATGCACTATGGCTGGCAAAGCGCACAC

NONO-PAM-mutation-fwd (C→G): ACCAGGAGAGAAGACGTTCACCCAACGAAGC

NONO-PAM-mutation-rev: GCTTCGTTGGGTGAACGTCTTCTCTCCTGGT

NONO-gateway-STOP-fwd (C→A): CAAACAAACGTCGCCGATACTAACCAACTTTCTTGTACAAAG

NONO-gateway-STOP-rev: CTTTGTACAAGAAAGTTGGTTAGTATCGGCGACGTTTGTTTG

**Plasmids for stable expression**

The desired pDONR 223-NONO construct was cloned into pLenti6.2-ccdB-3xFLAG-V5 (Addgene #87071) using the gateway vector conversion system (Invitrogen). Final plasmids contain a CMV promoter followed by a Gateway cloning linker, a start methionine, the NONO ORF without stop codon, a C-terminal FLAG tag, and a second Gateway cloning linker. All gene constructs were verified by DNA sequencing.

**NONO knockout by CRISPR/Cas9**

NONO knockout cells were generated using previously described protocols[12]. In brief, non-targeting control sgRNAs or sgRNAs targeting NONO (described below) were designed (https://portals.broadinstitute.org/gpp/public/analysis-tools/sgrna-design), and cloned into Lenti-CRISPR v2 plasmid (Addgene). 1 µg sgRNA-encoding plasmids were co-transfected with ΔVPR envelope (0.9 µg) and CMV VSV-G (0.1 µg) packaging plasmids into 2.5 × 106 HEK293T cells using the Fugene 6 transfection reagent (Promega). Virus-containing supernatants were collected forty-eight hours after transfection, passed through a 0.45 µM filter, and used to infect target cells in the presence of 10 µg/ml polybrene (Santa Cruz). Twenty-four hours post-infection, fresh media was added to the target cells, which were allowed to recover for an additional twenty-four hours. Puromycin (1 µg/mL) was then added to cells Following 10 days of puromycin selection, NONO expression in NONO sgRNA cells was determined by Western blot (see method below) compared to control sgRNA transfected cells.

Non-targeting sgRNA (sequences are 5′ to 3′):
Lenti-CRISPRv2: sgCRISPR-CTRL1-fwd: GCGAGGTATTCGGCTCCGCG
Lenti-CRISPRv2: sgCRISPR-CTRL1-rev: CGCGGAGCCGAATACCTCGC
sgRNAs targeting NONO (sequences are 5′ to 3′):
Lenti-CRISPRv2: NONO-1-fwd: CTGGACAATATGCCACTCCG
Lenti-CRISPRv2: NONO-1-rev: CGGAGTGGCATATTGTCCAG

**Generation of 22Rv1 NONO C-FLAG WT or C145S cell line**

1 µg pLenti6.2-NONO-C-FLAG-encoding plasmid (WT or C145S) with a silent PAM site mutation was co-transfected with ΔVPR envelope (0.9 µg) and CMV VSV-G (0.1 µg) packaging plasmids into 2.5 × $10^6$ HEK293T cells using the Fugene 6 transfection reagent (Promega). Virus-containing supernatants were collected forty-eight hours after transfection, passed through a 0.45 µM filter, and used to infect target cells in the presence of 10 µg/ml polybrene (Santa Cruz). Twenty-four hours post-infection, fresh media was added to the target cells, which were allowed to recover for an additional twenty-four hours. Blasticidin (10 µg/mL) was then added to cells. Following 10 days of blasticidin selection, NONO-FLAG expression was determined by Western blot (see method below) compared to empty vector transfected cells. Endogenous NONO was then knocked out by CRISPR/Cas9 using NONO sgRNA as described above.

**Western blot analysis**

For western blot protein analysis, cells (5x$10^5$ cells/treatment) were seeded in 2 mL media in a 6 well plate for 24h. Media was then aspirated and replaced with 2 mL media containing the indicated compounds or 0.1% DMSO for the indicated times. Following this incubation period, the cells were washed in cold DPBS, scraped, harvested and pelleted in 1.5 mL tubes (2000 g, 3 min, 4°C). Cell pellets were flash-frozen in liquid $N_2$ and stored at -80°C until further analysis. On the day of the analysis, the cell pellets were thawed on ice, re-suspended in cold PBS (100–150 µL), and lysed by sonication using a Branson Sonifier SFX250 (Branson Ultrasonics, catalog #101-063-965). Sonication was performed with a microtip probe at 10% amplitude, using pulsed mode (1 second on / 1 second off) for 2 cycles of 8 pulses each, while samples were kept on ice. Protein concentrations for all the samples were measured by BCA assay (ThermoFisher) and adjusted to 2 mg/mL, 4x loading buffer was added, and the samples were heated at 95°C for 5 min, followed by a 1 min spin at 20,000g. The proteins were resolved using SDS-PAGE (10% acrylamide gel) and transferred to 0.2 µm Power Blotter Select Transfer Stacks, nitrocellulose, mini (invitrogen). The membrane was blocked with 5% milk in Tris-buffered saline (20 mM Tris-HCl 7.6, 150 mM

NaCl) with 0.1% tween 20 (TBST) buffer at RT for 1 h or at 4°C overnight), washed with TBST, and incubated with primary antibodies in 5% milk in TBST at 4°C overnight (at 1:10000, 1:5000, or 1:2000 depending on the antibody). Following another TBST wash (3 x 3 min), the membrane was incubated with secondary antibody (1:1000 in 5% milk in TBST) at RT for 1h. The membrane was washed with TBST (3 x 3 min), developed with Femto (for RXRA, ESR1, or PSPC1) or standard (for NONO, FLAG, and GAPDH) ECL western blotting detection reagent kit (Thermo Scientific), and chemiluminescence was imaged on a ChemiDoc MP system (Bio-Rad). Relative band intensities were quantified using Image Lab software (Biorad).

**Gel-based ABPP**

5 x $10^5$ 22Rv1 cells stably expressing 3X-FLAG-V5 (WT or C145S), were seeded into a 6 well plate and allowed to adhere overnight at 37 °C. Once at 80-90% confluence, cells were then treated with DMSO (0.1%) or the indicated compound for 6h *in situ*, followed by treatment with (*R*, *R*)-GL-586 or #14 alkyne at the indicated concentration (otherwise 10 μM) probe *in situ* for 1h at 37 °C. Following this incubation period, the cells were washed in cold DPBS, scraped, harvested and pelleted in 1.5 mL tubes (2000 g, 5 min, 4°C). Cell pellets were flash-frozen in liquid $N_2$ and stored at -80°C until further analysis. On the day of the analysis, the cell pellets were thawed on ice, re-suspended in cold PBS (with Roche cOmplete, mini, EDTA-free Protease Inhibitor Cocktail, 1 tablet in 10 mL PBS) and lysed by sonication with Branson probe sonicator (2 x 15 pulses; 10% power output). Protein concentrations for all the samples were measured by BCA assay (Thermofischer) and adjusted to 2 mg/mL. Rhodamine-azide (1 μL/reaction, 1.25 mM in DMSO), $CuSO_4$ (1 μL/reaction, 50 mM in $H_2O$), TBTA (3 μL/reaction, 1.7 mM in DMSO/t-BuOH (1:4, v/v)) and tris(2-carboxyethyl) phosphine (TCEP) (1 μL/reaction, 50 mM in $H_2O$, freshly prepared) were premixed. 6 μL of this click reagent mixture was immediately added to 50 μL of each alkyne probe-labeled sample and incubated for 1 h at room temperature. The reactions were quenched by adding 4X SDS–PAGE loading buffer. The quenched samples were loaded on a 10% acrylamide gel for separation by SDS-PAGE. Samples were visualized by in-gel fluorescence scanning using the ChemiDoc MP system (Bio-Rad) and fluorescent band intensity was quantified with Image Lab software (Bio-Rad).

**Activity-Based Protein Profiling (ABPP) Methods**

**Cysteine-directed ABPP**

Cysteine-directed activity based-protein profiling (ABPP) was performed as previously described.[13] 22Rv1 cells (2 million cells in 10 cm dish) were plated and given time to adhere to the plate. Once plates were about 80-90% confluent the cells were treated with DMSO or indicated probes. Cells were washed with ice-cold DPBS (3x), followed by resuspension in DPBS and lysed by probe-sonication (2 x 15 pulses; 10% power output). The total protein content of whole-cell lysates was measured using a Pierce BCA protein assay kit and the samples were normalized to 2 mg/mL and 500 µL. Samples were treated with 5 µL of 10 mM IA-DTB (in DMSO) for 1 h at RT with vortexing every 20 min. Proteins were precipitated by the addition of cold methanol (600 µL), chloroform (200 µL) and HPLC-grade water (100 µL), followed by vortexing and centrifugation at 16,000 x g for 10 min. Without disrupting the protein disk, both the top and bottom layers were aspirated, and the protein disk was sonicated again in 500 µL of methanol and centrifuged at 16,000 x g for 10 min. After the methanol was completely aspirated, protein pellets were immediately processed or frozen at −80°C. Pellets were resuspended in 90 µL of denaturing/reducing buffer (9 M urea, 10 mM DTT, 50 mM triethylammonium bicarbonate (TEAB) pH 8.5). The samples were reduced by heating at 65°C for 20 min, followed by alkylation with 10 µL of 500 mM iodoacetamide at 37°C for 30 min. The samples were then centrifuged at 16,000 x g for 2 min to pellet any insoluble precipitate and probe-sonicated once more to ensure complete resuspension, and then diluted with 300 µL of 50 mM TEAB pH 8.5 to reach a final urea concentration of 2 M. Trypsin (4 µL of 0.25 µg/µL in trypsin resuspension buffer with 25 mM $CaCl_2$) was added to each sample and digested at 37 °C overnight. Digested samples were then diluted with 300 µL of enrichment buffer (50 mM TEAB pH 8.5, 150 mM NaCl, 0.2% NP-40) containing streptavidin-agarose beads (50 µL of 50% slurry/sample) and were rotated at RT for 2 h. The samples were centrifuged (2,000 x g, 2 min) and the entire content transferred to BioSpin columns and washed (3 x 1 mL wash buffer, 3 x 1 mL DPBS, 3 x 1 mL water). Enriched peptides were eluted from the beads with 300 µL of 50% acetonitrile with 0.1% formic acid and dried using a SpeedVac at 46°C. Enriched peptides were resuspended in 100 µL EPPS buffer (200 mM, pH 8.0) with 30% acetonitrile, vortexed and water bathsonicated. The samples were TMT-labelled by the addition of 3 µL of 20 mg/mL (in dry acetonitrile) of corresponding $TMT^{10}$plex tag for 1.5 h at RT with vortexing every 30 min. TMT labelling was quenched by the addition of hydroxylamine (3 µL 5% solution in $H_2O$) and incubated for 15 min at RT. Samples were then acidified with 5 µL formic acid, combined and dried using a SpeedVac. Samples were desalted with a Sep-Pak column and then high-pH-fractionated by HPLC (described in the following section) into a 96-well plate and recombined into 12 fractions (total).

**HPLC fractionation**

Desalted samples were resuspended in 500 mL buffer A (5% acetonitrile, 0.1% formic acid in milliQ water) and fractionated with Agilent HPLC into a 96 deep-well plate containing 20 mL of 20% formic acid to acidify the eluting peptides, as previously reported.28 The peptides were eluted onto a capillary column (ZORBAX 300Extend-C18, 3.5 mm) and separated at a flow rate of 0.5 mL/min using the following gradient: 100% buffer A from 0-2 min, 0%–13% buffer B from 2-3 min, 13%–42% buffer B from 3-60 min, 42%–100% buffer B from 60-61 min, 100% buffer B from 61-65 min, 100%–0% buffer B from 65-66 min, 100% buffer A from 66-75 min, 0%–13% buffer B from 75-78 min, 13%–80% buffer B from 78-80 min, 80% buffer B from 80-85 min, 100% buffer A from 86-91 min, 0%–13% buffer B from 91-94 min, 13%–80% buffer B from 94-96 min, 80% buffer B from 96-101 min, and 80%–0% buffer B from 101-102 min (buffer A: 10 mM aqueous NH4HCO3; buffer B: acetonitrile). The plates were evaporated to dryness using SpeedVac and peptides resuspended in 80% acetonitrile, with 0.1% formic acid and combined to a total of 12 fractions (e.g., fraction1= well 1A+ 1B.1H, fraction 2= well 2A+2B..2H) (3x300 mL/column). Samples were SpeedVac to dryness and the resulting 12 fractions were re-suspended in buffer A (5% acetonitrile, 0.1% formic acid) and analyzed by mass spectrometry.

**Protein-directed ABPP**

Protein-directed ABPP was performed as previously described.[13] MCF7 cells were treated with DMSO or an indicated concentration of compounds for 6 hours (unless otherwise stated) followed by an alkyne probe (10 µM, 1h). Cells were washed with ice-cold DPBS (3x). Cell pellets were resuspended in DPBS and lysed by probe-sonication (2x15 pulses; 10% power output). Proteome was normalized to 2 mg/mL in 500 mL (Pierce BCA protein assay), and the alkyne probe labeled proteins were treated with 55 µL of click MS-Master-mix [30 µL of 1.7 mM TBTA in 4:1 t-BuOH:DMSO, 10 µL of 50 mM $CuSO_4$ in H2O, 10 µL of freshly prepared 50 mM Tris(2-carboxyethyl)phosphine in H2O, 10 µL of 10 mM Biotin-PEG4-azide]. Proteins were precipitated with cold methanol (600 µL), chloroform (200 µL) and water (100 µL), vortexed, and then centrifuged at 16,000 x g for 10 min. The top and bottom layers were aspirated, and the protein-disk was washed with 1 mL ice-cold MeOH and the protein disk was then sonicated in 500 µL of ice-cold methanol and pelleted at 16,000 x g for 10 min. After the methanol was completely aspirated, protein pellets were immediately processed or stored at −80°C. Pellets were resuspended in 500 µL freshly prepared 8 M urea in DPBS, followed by the addition of 10 µL of 10 wt% SDS. Samples were then pulse-sonicated until clear (~15 pulses). The samples were reduced with 25 µL of 200 mM dithiothreitol (DTT) at 65°C for 15 min, followed by alkylation with 25 µL of 400 mM iodoacetamide at 37°C for 30 min in the dark. Then, 65 µL of 20 wt% SDS was added, and the samples were transferred to a 15-mL tube in a total volume of 6 mL with DPBS (0.2% final SDS). Washed streptavidin beads (Thermo, #20353; 100 µL of 50% slurry/sample) were then added and proteins were enriched for 1.5 h at RT with rotation. After incubation, the beads were pelleted (2 min at 2,000 x g) and washed with 0.2% wt% SDS in DPBS (2 x 10 mL), DPBS (1 x 5 mL), HPLC-grade water (2 x 1 mL) and 200 mM 4-(2- hydroxyethyl)-1-piperazinepropanesulfonic acid (EPPS; 1 mL, pH 8.0). Enriched proteins were digested on-bead overnight with 200 µL of trypsin mix (2 M urea, 1 mM CaCl2, 10 µg/mL trypsin (Promega, #V5111), 200 mM EPPS, pH 8.0). The beads were pelleted at 2,000 x g, the supernatant was collected and then diluted with 100 µL acetonitrile (30% final). Samples were then labeled with 6 µL 20 mg/mL (in dry acetonitrile) TMTpro$^{TM}$ 16plex tag (Thermo, #A44520) or TMT10plex$^{TM}$ (Thermo, #90406) for 1.5 h at RT (vortexed every 30 min). TMT labelling was quenched by the addition of hydroxylamine (6 µL 5% solution in $H_2O$) and incubated for 15 min at RT. Samples were then acidified with 20 µL formic acid, combined and dried using a SpeedVac at 46 °C. Samples were desalted with a Sep-Pak column Vac 1 cc (50 mg) (Waters, #WAT054955) and then high pH fractionated into ten fractions using Pierce peptide desalting spin columns (Thermo, #89852) and an acetonitrile/NH4HCO3 (10 mM) gradient for high-pH spin column fractionation (described in the following section) and analyzed by mass spectrometry.

**High-pH spin column fractionation**

Thirty high-pH elution buffers were freshly prepared for peptide fractionation using increasing concentrations of acetonitrile (ACN) in 10 mM aqueous ammonium bicarbonate ($NH_4HCO_3$). Each eluent was prepared in low-bind microcentrifuge tubes by mixing the appropriate volumes of ACN and 10 mM $NH_4HCO_3$ to a final volume of 1 mL per fraction. The ACN concentrations ranged from 7.5% to 95%, increasing in 2.5% increments. For each sample, one high-pH reversed-phase spin column was equilibrated. To begin, the white bottom cap of the spin column was removed, and the red top cap was loosely tightened. The spin column was placed into a 2.0 mL microcentrifuge tube and centrifuged at 5,000 × g for 2 minutes to pack the resin and remove the storage solution. Next, each column was washed twice with 300 µL of 100% acetonitrile (MeCN), centrifuging at 5,000 × g for 2 minutes each time and discarding the flow-through. This was followed by two washes with 300 µL of water containing 0.1% formic acid, using the same centrifugation conditions. The column was then considered equilibrated. Dried, TMT-labeled peptide samples were resuspended in 300 µL of buffer A and loaded onto the equilibrated column. The sample was centrifuged at 2,000 × g for 2 minutes, and the flow-through was collected, re-applied to the column, and centrifuged again at 2,000 × g for 2 minutes to maximize peptide retention. The column was washed by transferring it to a fresh 2.0 mL microcentrifuge tube and adding 300 µL of water, followed by centrifugation at 2,000 × g for 2 minutes. To remove excess TMT reagent, 300 µL of 5% MeCN in 10 mM aqueous ammonium bicarbonate ($NH_4HCO_3$) was added to the column, and the sample was centrifuged at 2,000 × g for 2 minutes. Peptides were eluted into fresh 1.5 mL microcentrifuge tubes using the pre-prepared high-pH eluent mixtures (100 µL per fraction, 300 µL for the final elution), spinning at 2,000 × g for 1 minute per elution. A total of 30 individual fractions were collected. Every 10th fraction was combined to yield 10 final pooled fractions (e.g., fractions 1, 11, and 21 were combined). Combined fractions were dried using a SpeedVac concentrator and analyzed by mass spectrometry.

**TMT LC-MS Analysis**

Fractions were resuspended in buffer A (5% acetonitrile, 0.1% formic acid in water) and analyzed by liquid chromatography tandem mass-spectrometry using an Orbitrap Fusion Tribrid Mass Spectrometer (Thermo Scientific) coupled to an UltiMate 3000 Series Rapid Separation LC system and autosampler (Thermo Scientific Dionex). The peptides were eluted onto a capillary column (75-µm-inner-diameter fused silica, packed with C18 (Waters, Acquity BEH C18, 1.7 µm, 25 cm) or an EASY-Spray HPLC column (Thermo, #ES902, #ES903) using an Acclaim PepMap 100 (Thermo, #164535) loading column, and separated at a flow rate of 0.25 µL min-1. Peptides were separated across a 10 min gradient of 5%, 150 min gradient of 5-20%, 20 min 20-45%, and then 5 min 45-95% acetonitrile (0.1% formic acid) in H2O (0.1% formic acid) followed by column equilibration. Data were acquired using an MS3-based TMT method on Orbitrap Fusion or Eclipse Tribrid mass spectrometers.

Fusion instruments: The scan sequence began with an MS1 master scan (Orbitrap analysis, resolution 120,000, 400-1,700 m/z, RF lens 60%, maximum injection time 50 ms) with dynamic exclusion enabled (repeat count 1, duration 15 s). The top precursors were then selected for MS2/MS3 analysis. MS2 analysis consisted of quadrupole isolation (isolation window 0.7) of precursor ion followed by collision-induced dissociation in the ion trap (collision energy 35%, maximum injection time 120 ms). Following the acquisition of each MS2 spectrum, synchronous precursor selection enabled the selection of up to 10 MS2 fragment ions for MS3 analysis. MS3 precursors were fragmented by higher energy collisional dissociation (HCD) and analyzed using the Orbitrap (collision energy 55, maximum injection time 120 ms, resolution 50,000). For MS3 analysis, we used charge state-dependent isolation windows. For charge state z = 2, the MS isolation window was set at 1.2; for z = 3–6 the MS isolation window was set at 0.7.

Eclipse Tribrid instrument: The scan sequence began with an MS1 master scan (Orbitrap analysis, resolution 120,000, 400-1,700 m/z, RF lens 30%, maximum injection time 50 ms) with dynamic exclusion enabled (repeat count 1, duration 30 s). The top precursors were then selected for MS2/MS3 analysis. MS2 analysis consisted of quadrupole isolation (isolation window 0.7) of precursor ion followed by higher-energy collisional dissociation (HCD) in the ion trap (collision energy 36%, maximum injection time 120 ms). Following the acquisition of each MS2 spectrum, synchronous precursor selection enabled the selection of up to 10 MS2 fragment ions for MS3 analysis. MS3 precursors were fragmented by HCD and analyzed using the Orbitrap (collision energy 55%, maximum injection time 120 ms, resolution 30,000). For MS3 analysis, we used charge state dependent isolation windows. For charge state $z = 2$, the MS isolation window was set at 1.2; for $z = 3$ the MS isolation window was set at 0.7; for $z = 4$–6, the MS isolation window was set at 0.4.

**MS data processing**

Raw files were uploaded to the Integrated Proteomics Pipeline (IP2, version 6.7.1) available at http://ip2.scripps.edu/ip2/mainMenu.html and MS2 and MS3 files were extracted from the raw files using RAW Converter (version 1.1.0.22) available at http://fields.scripps.edu/rawconv/ and searched using the ProLuCID algorithm using a reverse concatenated, non-redundant variant of the Human UniProt database (release 2016-07). Cysteine residues were searched with a static modification for carboxyamidomethylation (+57.02146 Da). N-termini and lysine residues were also searched with a static modification corresponding to the TMT tag (+229.1629 Da for 10-plex and +304.2071 Da for 16-plex). Peptides were required to be at least six amino acids long. ProLuCID data were filtered through DTASelect (version 2.0) to achieve a spectrum false-positive rate below 1%. We included a keratin filter. The MS3-based peptide quantification was performed with reporter ion mass tolerance set to 20 ppm with the Integrated Proteomics Pipeline (IP2).

**Data analysis – Cysteine-directed ABPP**

The census output files from Integrated Proteomics Pipeline 2 (IP2, v.6.7.1) were further processed to calculate cysteine engagement ratios (probe vs DMSO) by dividing each TMT reporter ion intensity by the average intensity for the DMSO channels. Peptide-spectra matches were grouped based on protein ID and the cysteine residue number. Peptides with summed reporter ion intensities < 10000, coefficient of variation for DMSO channels > 0.5 were excluded from analysis. TMT ion intensities were median normalized per TMT channel. Two independent replicates of 10-plex experiments in 22Rv1 cells were analyzed together (total n = 2-4 per treatment condition). Data were averaged from all replicates.

**Data analysis– Protein-directed ABPP**

The census output files from IP2 were further processed to calculate enrichment engagement ratios (probe vs probe) by dividing each TMT reporter ion intensity by the sum of intensity for all the channels. Each spectrum-peptide match was grouped based on protein ID, excluding peptides with summed reporter ion intensities < 10,000, coefficient of variation of > 0.5, < 2 unique peptides per protein ID.

**RNA-Seq**

**Sample preparation**

MCF7 cells (1x$10^6$ cells/treatment) were seeded in 2 mL media in a 6 well cm plate for 24h. Media was then aspirated and replaced with 2 mL media containing the indicated compounds or 0.1% DMSO for 6h. Following this incubation period, the cells were washed in cold DPBS, scraped, harvested and pelleted in 1.5 mL tubes (2000 g, 3 min, 4°C). Cell pellets were flash-frozen in liquid $N_2$ and stored at -80°C until further analysis. Total RNA from thawed cells was isolated using RNeasy Plus Kit (QIAGEN) with QIAshredder columns for cell lysis (QIAGEN) according to the manufacturer's protocol and stored at -80°C until further analysis. RNA concentration was measured by Nanodrop and 1-2 µg were used for sequencing.

**Library preparation with polyA selection and HiSeq Sequencing**

Library preparations and sequencing reactions were conducted at GENEWIZ, LLC. (South Plainfield, NJ, USA) as follows: Extracted RNA samples were quantified using Qubit 2.0 Fluorometer (Life Technologies) and RNA integrity was checked using Agilent TapeStation 4200 (Agilent Technologies). RNA sequencing libraries were prepared using the NEBNext Ultra RNA Library Prep Kit for Illumina following manufacturer's instructions (NEB). Briefly, mRNAs were first enriched with Oligo(dT) beads. Enriched mRNAs were fragmented for 15 minutes at 94 °C. First strand and second strand cDNAs were subsequently synthesized. cDNA fragments were end repaired and adenylated at 3' ends, and universal adapters were ligated to cDNA fragments, followed by index addition and library enrichment by limited-cycle PCR. The sequencing libraries were validated on the Agilent TapeStation (Agilent Technologies) and quantified by using Qubit 2.0 Fluorometer (Invitrogen) as well as by quantitative PCR (KAPA Biosystems). The sequencing libraries were clustered on 2 lanes of a flowcell. After clustering, the flowcell was loaded on the

Illumina HiSeq instrument (4000 or equivalent) according to manufacturer's instructions. The samples were sequenced using a 2x150bp Paired End (PE) configuration. Image analysis and base calling were conducted by the HiSeq Control Software (HCS). Raw sequence data (.bcl files) generated from Illumina HiSeq was converted into fastq files and de-multiplexed using Illumina's bcl2fastq 2.17 software. One mismatch was allowed for index sequence identification.

**RNA-seq data processing and quantification**

Raw RNA-seq reads were processed using the nf-core/rnaseq pipeline (v3.13.2),[14] which includes quality control with FastQC, adapter trimming, and alignment to the human reference genome (GRCh38) using STAR Salmon. Aligned reads were quantified at the gene level using *featureCounts* (v2.0.6) from the Subread package, with annotations derived from GENCODE release v29. Default parameters were used, and count matrices were generated for downstream differential expression analysis.

***RNA-seq data analysis***

Raw read counts were generated using FeatureCounts (v2.0.1) from aligned BAM files. Differential expression analysis was performed using the DESeq2 package (v1.40.1) in R (v4.3.1). Samples were initially collected in biological triplicate for each condition. However, quality control analysis revealed that two samples in different treatment groups behaved as outliers. These replicates were removed from the analysis to avoid skewing downstream results. To avoid unbalanced statistics, a principal component analysis based on variance-stabilizing transformation (VST) was used to remove one replicate in each group of triplicates that clustered separately from the other two. Final differential expression analysis was therefore performed using two biological replicates per condition. DESeq2's standard pipeline was used, including size factor estimation, dispersion estimation, and negative binomial model fitting. Wald tests were used to determine statistical significance, and Benjamini-Hochberg correction was applied to control the false discovery rate (FDR). Genes with an adjusted p-value < 0.01 were considered significantly differentially expressed.

**Cell growth inhibition assays**

Cells were seeded at 5000 cells per well (100 µL) in 96-well clear-bottom, white-wall plates. 100 µL of medium containing DMSO or compounds (2 x final concentrations) were added and cultured for 5 days (replacing media + compound every 48 hours). On the day of reading, media was removed and replaced with 100 µL of fresh media before adding 100 µL CellTiterGlo reagent (Promega) for 30 minutes. Luminescence was then measured on a Clariostar plate reader (BMG Labtech). Relative cell growth was determined by normalizing the luminescence reading to the DMSO treated control.

**Quantification and statistical analysis**

Statistical analysis was performed using GraphPad Prism version 9 for Windows (GraphPad Software, La Jolla California USA, https://www.graphpad.com/). Statistical values including the n and statistical significance are reported in the Figure legends. All details are included in figure legends.

### Supplementary Information: Synthesis and Characterization of Novel Compounds

### General considerations

All NMR spectra were recorded at 298 K unless otherwise noted. $^{1}$H NMR spectra were recorded on Bruker Avance III 400, Avance III HD 400, Avance Neo 400 spectrometers ($^{1}$H, 400 MHz). $^{13}$C NMR spectra were recorded on a ZKNJ QOne Quantum-I Plus 400 spectrometer ($^{13}$C, 101 MHz). $^{19}$F NMR spectra were recorded on a Bruker AV Neo 399 MHz spectrometer ($^{19}$F, 376 MHz). $^{1}$H NMR data are reported as follows: chemical shift (δ), multiplicity (s = singlet, d = doublet, t = triplet, m = multiplet; br = broad), coupling constants, and integration. Chemical shifts are reported in parts per million (ppm) using the appropriate solvent as reference.[15] Analytical supercritical fluid chromatography (SFC) was performed on a Shimadzu LC system (flow rate: 3 mL/min, back pressure: 100 Bar, column temperature: 35 °C) equipped with a polydiode array detector. Mass measurements for high-resolution mass spectrometry (HRMS) were performed on a Waters Xevo G2-XS TOF calibrated against sodium formate clusters and using a LeuEnk lockmass. Expected monoisotopic masses were calculated using MassLynx 4.1 and the m/z values for calibrant and lockmass were MassLynx-default values.

### Synthesis of (*R*,*R*)-GL-373

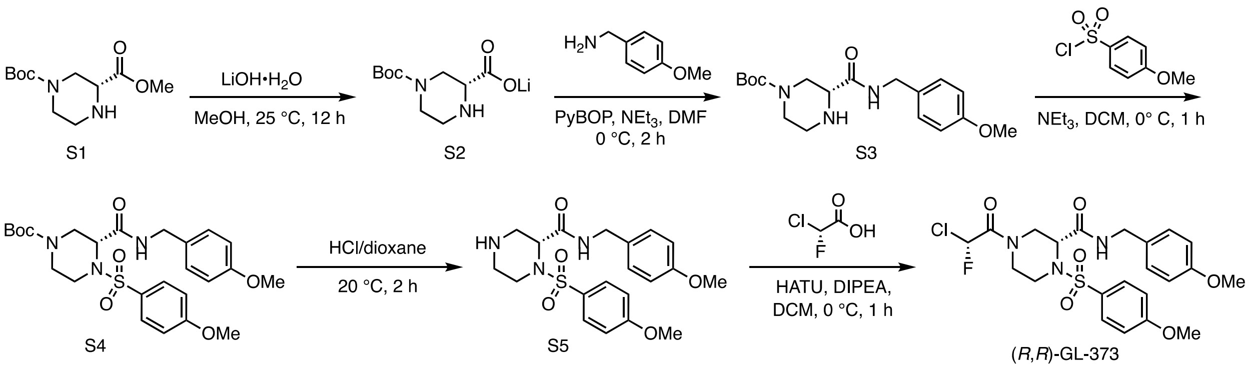


**Figure S5.** Synthesis of (*R*,*R*)-GL-373.

#### Lithium (*R*)-4-(*tert*-butoxycarbonyl)piperazine-2-carboxylate (S2)

To a solution of **S1** (2.00 g, 8.19 mmol, 1.0 equiv) in MeOH (20 mL) were added LiOH•$H_2O$ (515 mg, 12.3 mmol, 1.5 equiv) and $H_2O$ (2 mL). The mixture was stirred at 25 °C for 12 h. On completion, the reaction mixture was concentrated under reduced pressure to give **S2** (1.95 g, quant.) as a yellow solid, which was used in the next step without further purification.

**tert-butyl (R)-3-((4-methoxybenzyl)carbamoyl)piperazine-1-carboxylate (S3)**

To a precooled (0 ºC) solution of **S2** (300 mg, 1.27 mmol, 1.0 equiv) and 4-methoxybenzylamine (192 mg, 1.40 mmol, 1.1 equiv) in DMF (5 mL) were added PyBOP (991 mg, 1.91 mmol, 1.5 equiv) and triethylamine (386 mg, 3.81 mmol, 3.0 equiv). The mixture was stirred at 0 °C for 2 h. On completion, the reaction mixture was concentrated under reduced pressure to give a residue, which was purified by reverse-phase-HPLC (A: $H_2O$ (10mM $NH_4HCO_3$), B: acetonitrile; gradient: B%: 0%-100%, 20 min) and purified by prep-HPLC (column: Waters Xbridge Prep OBD C18 150*40mm*10um; mobile phase: [$H_2O$ (0.05% $NH_3$•$H_2O$)-acetonitrile]; gradient: 29%-59% B over 15.0 min) to give **S3** (0.30 g, 68% yield) as a white solid.

$^1$H NMR (400 MHz, $CDCl_3$) δ 7.19 (d, *J* = 8.2 Hz, 2H), 7.11 – 6.94 (m, 1H), 6.86 (d, *J* = 8.8 Hz, 2H), 4.37 (d, *J* = 5.8 Hz, 2H), 4.14 – 4.06 (m, 1H), 3.84 – 3.73 (m, 1H), 3.80 (s, 3H), 3.35 (dd, *J* = 9.3, 3.6 Hz, 1H), 3.01 (t, *J* = 11.4 Hz, 1H), 2.91 (d, *J* = 12.3 Hz, 2H), 2.76 (t, *J* = 11.4 Hz, 1H), 1.45 (s, 9H), 1 exchangeable proton not observed.

LC-MS m/z calc. for $C_{18}H_{28}N_3O_4$ $[M+H]^+$ 350.2 found 350.2.

**tert-butyl (R)-3-((4-methoxybenzyl)carbamoyl)-4-((4-methoxyphenyl)sulfonyl)piperazine-1-carboxylate (S4)**

To a precooled (0 ºC) solution of **S3** (0.30 g, 859 µmol, 1.0 equiv) and triethylamine (261 mg, 2.58 mmol, 3.0 equiv) in dichloromethane (10 mL) was added (4-methoxyphenyl)sulfonyl chloride (266 mg, 1.29 mmol, 1.5 equiv). The mixture was stirred at 0 °C for 2 h. On completion, the reaction mixture was concentrated under reduced pressure. The resulting residue was purified by flash column chromatography ($SiO_2$, petroleum ether/EtOAc = 5:1 to 2:1) to give **S4** (0.35 g, 78% yield) as a white solid.

$^1$H NMR (400 MHz, $CDCl_3$) δ 7.75 (d, *J* = 9.0 Hz, 2H), 7.15 (d, *J* = 8.6 Hz, 2H), 6.97 (d, *J* = 9.0 Hz, 2H), 6.85 (d, *J* = 8.7 Hz, 2H), 6.82 – 6.76 (m, 1H), 4.63 – 4.51 (m, 1H), 4.48 – 4.25 (m, 3H), 3.87 (s, 3H), 3.80 (s, 3H), 3.80 – 3.57 (m, 2H), 3.32 – 3.21 (m, 1H), 2.91 – 2.58 (m, 2H), 1.41 (s, 9H).

LC-MS m/z calc. for $C_{25}H_{34}N_3O_7S$ $[M+H]^+$ 520.2 found 520.2.

**(R)-N-(4-methoxybenzyl)-1-((4-methoxyphenyl)sulfonyl)piperazine-2-carboxamide (S5)**

To a solution of **S4** (1.00 g, 1.92 mmol) in dioxane (2 mL) was added HCl (2 M solution in dioxane, 5 mL, 5.2 equiv).The mixture was stirred at 20 °C for 2 h. On completion, the reaction mixture was concentrated in vacuo to give **S5•HCl** (750 mg, 90% w/w [dioxane], 77% yield) as a white solid, which was used in the next step without further purification.

$^1$H NMR (400 MHz, $CD_3OD$) δ 8.45 (t, *J* = 4.4 Hz, 1H), 7.80 (d, *J* = 8.9 Hz, 2H), 7.19 (d, *J* = 8.7 Hz, 2H), 7.04 (d, *J* = 9.0 Hz, 2H), 6.88 (d, *J* = 8.7 Hz, 2H), 4.76 (d, *J* = 4.5 Hz, 1H), 4.33 – 4.15 (m, 2H), 4.00 (dd, *J* = 14.5, 3.8 Hz, 1H), 3.88 (s, 3H), 3.79 (s, 3H), 3.72 – 3.59 (m, 3H + dioxane), 3.34 (m, 1H), 3.07 (dd, *J* = 13.2, 4.6 Hz, 1H), 2.96 (td, *J* = 12.8, 4.1 Hz, 1H).

LC-MS m/z calc. for $C_{20}H_{26}N_3O_5S$ $[M+H]^+$ 420.2 found 420.2.

#### (*R*)-4-((*R*)-2-chloro-2-fluoroacetyl)-*N*-(4-methoxybenzyl)-1-((4-methoxyphenyl)sulfonyl) piperazine-2-carboxamide ((*R,R*)-GL-373)

To a precooled (0 ºC) solution of **S5•HCl** (160 mg, 90% w/w [dioxane], 316 μmol, 1.0 equiv), DIPEA (181 mg, 1.40 mmol, 4.4 equiv) and (*R*)-chlorofluoroacetic acid (59.2 mg, 526 μmol, 1.7 equiv) in dichloromethane (2 mL) was added HATU (267 mg, 702 μmol, 2.2 equiv). The mixture was stirred at 0 °C for 1 h. On completion, the reaction mixture was concentrated in vacuo. The residue was purified by prep-TLC ($SiO_2$, petroleum ether/EtOAc = 2:1) and prep-HPLC (column: Waters xbridge 150*25mm 10μm; mobile phase: A: $H_2O$ (10mM $NH_4HCO_3$), B: acetonitrile; gradient: 30%-60% B over 15.0 min) to give **(*R*,*R*)-GL-373** (80.0 mg, 49% yield) as a white amorphous solid.

Note: NMR spectra are consistent with a 7:3 mixture of rotamers.

$^1$H NMR (400MHz, $CDCl_3$): δ 7.73 (d, *J* = 9.0 Hz, 2H), 7.20 – 7.07 (m, 2H), 7.04 – 6.93 (m, 3H), 6.92 – 6.76 (m, 2.7H), 6.35 (d, $^2J_{HF}$ = 50.4 Hz, 0.3H), 4.84 (d, *J* = 13.7 Hz, 0.3H), 4.56 – 4.41 (m, 1.7H), 4.41 – 4.19 (m, 2H), 4.11 (d, *J* = 13.7 Hz, 0.7H), 3.99 – 3.65 (m, 7.3H), 3.55 – 3.38 (m, 0.3H), 3.31 – 3.07 (m, 1H), 2.99 (dd, *J* = 14.1, 4.2 Hz, 0.7H), 2.77 (dd, *J* = 14.0, 4.5 Hz, 0.3H), 2.65 (t, *J* = 12.3 Hz, 0.7H).

$^{19}$F NMR (376 MHz, $CDCl_3$): δ 142.93, 139.69.

$^{13}$C NMR (101 MHz, $CDCl_3$) δ 167.32, 167.18, 163.79, 163.59, 161.90, 161.68, 159.14, 130.11, 129.41, 129.30, 129.12, 128.93, 115.04, 114.75, 114.21, 114.08, 97.15, 95.48, 94.69, 92.91, 56.03, 55.77, 55.73, 55.30, 50.79, 43.59, 43.54, 43.44, 43.25, 42.50, 42.25, 41.95, 41.12.

HRMS m/z calc. for $C_{22}H_{26}ClFN_3O_6S$ $[M+H]^+$ 514.1215 found 514.1217.

### Synthesis of stereoisomers of (*R*,*R*)-GL-373

#### (*S*)-4-((*S*)-2-chloro-2-fluoroacetyl)-*N*-(4-methoxybenzyl)-1-((4-methoxyphenyl)sulfonyl) piperazine-2-carboxamide ((*S*,*S*)-GL-373)

**(*S*,*S*)-GL-373** was prepared from *ent*-**S5•HCl** and (*S*)-chlorofluoroacetic acid following a procedure analogous to that used for the synthesis of (*R*,*R*)-GL-373.

Note: NMR spectra are consistent with a 7:3 mixture of rotamers.

$^1$H NMR (400 MHz, $CDCl_3$) δ 7.74 (d, *J* = 8.9 Hz, 2H), 7.18 – 7.09 (m, 2H), 7.06 – 6.89 (m, 3.3H), 6.88 – 6.75 (m, 2.4H), 6.35 (d, $^2J_{HF}$ = 50.4 Hz, 0.3H), 4.85 (d, *J* = 13.7 Hz, 0.3H), 4.56 – 4.42 (m, 1.7H), 4.42 – 4.20 (m, 2H), 4.12 (d, *J* = 13.7 Hz, 0.7H), 4.00 – 3.67 (m, 7.3H), 3.45 (t, *J* = 12.3 Hz, 0.3H), 3.30 – 3.09 (m, 1H), 2.99 (dd, *J* = 14.0, 4.2 Hz, 0.7H), 2.85 – 2.73 (m, 0.3H), 2.65 (t, *J* = 12.3 Hz, 0.7H).

$^{19}F$ NMR (376 MHz, $CDCl_3$): δ 142.84, 139.64.

$^{13}C$ NMR (101 MHz, $CDCl_3$) δ 167.28, 167.09, 163.81, 161.89, 161.67, 159.16, 130.12, 129.87, 129.33, 129.16, 128.95, 115.06, 114.77, 114.22, 97.22, 94.82, 56.07, 55.82, 55.39, 55.27, 43.49, 42.51, 41.84, 41.10.

HRMS m/z calc. for $C_{22}H_{26}ClFN_3O_6S$ $[M+H]^+$ 514.1215 found 514.1219.

**(*R*)-4-((*S*)-2-chloro-2-fluoroacetyl)-*N*-(4-methoxybenzyl)-1-((4-methoxyphenyl)sulfonyl) piperazine-2-carboxamide ((*S*,*R*)-GL-373)**

**(*S*,*R*)-GL-373** was prepared from **S5•HCl** and (*S*)-chlorofluoroacetic acid following a procedure analogous to that used for the synthesis of (*R*,*R*)-GL-373.

Note: NMR spectra are consistent with a 9:1 mixture of rotamers.

$^{1}H$ NMR (400 MHz, $CDCl_3$) δ 7.75 (d, $J$ = 9.0 Hz, 2H), 7.17 – 6.93 (m, 6H), 6.89 – 6.81 (m, 2H), 6.34 (d, $^{2}J_{HF}$ = 50.7 Hz, 0.1H), 4.82 (d, $J$ = 13.7 Hz, 0.1H), 4.45 (d, $J$ = 3.7 Hz, 1H), 4.41 – 4.20 (m, 4H), 3.95 – 3.83 (m, 4H), 3.79 (s, 3H), 3.11 (ddd, $J$ = 15.1, 12.1, 3.4 Hz, 1H), 2.81 (dd, $J$ = 14.0, 4.0 Hz, 1H), 2.45 (td, $J$ = 13.1, 3.5 Hz, 1H).

$^{19}F$ NMR (376 MHz, $CDCl_3$): δ 147.73, 141.33.

$^{13}C$ NMR (101 MHz, $CDCl_3$) δ 167.23, 167.13, 163.83, 162.95, 162.70, 159.19, 159.08, 130.47, 129.93, 129.26, 129.21, 129.08, 128.90, 115.15, 114.83, 114.27, 91.11, 91.04, 88.64, 88.59, 56.06, 56.01, 55.89, 55.74, 55.40, 55.26, 43.56, 43.41, 42.86, 42.77, 40.42, 40.32.

HRMS m/z calc. for $C_{22}H_{26}ClFN_3O_6S$ $[M+H]^+$ 514.1215 found 514.1209.

**(*S*)-4-((*R*)-2-chloro-2-fluoroacetyl)-*N*-(4-methoxybenzyl)-1-((4-methoxyphenyl)sulfonyl) piperazine-2-carboxamide ((*R*,*S*)-GL-373)**

**(*R*,*S*)-GL-373** was prepared from *ent*-**S5•HCl** and (*R*)-chlorofluoroacetic acid following a procedure analogous to that used for the synthesis of (*R*,*R*)-GL-373.

Note: NMR spectra are consistent with a 9:1 mixture of rotamers.

$^{1}H$ NMR (400 MHz, $CDCl_3$) δ 7.75 (d, $J$ = 8.9 Hz, 2H), 7.17 – 6.93 (m, 6H), 6.84 (d, $J$ = 8.6 Hz, 2H), 6.35 (d, $^{2}J_{HF}$ = 50.6 Hz, 0.1H), 4.82 (d, $J$ = 13.7 Hz, 0.1H), 4.53 – 4.43 (m, 1H), 4.41 – 4.18 (m, 4H), 3.94 – 3.83 (m, 4H), 3.79 (s, 3H), 3.11 (ddd, $J$ = 15.0, 12.1, 3.4 Hz, 1H), 2.81 (dd, $J$ = 14.0, 4.0 Hz, 1H), 2.45 (td, $J$ = 12.8, 3.5 Hz, 1H).

$^{19}F$ NMR (376 MHz, $CDCl_3$): δ 147.73, 141.34.

$^{13}C$ NMR (101 MHz, $CDCl_3$) δ 167.23, 167.14, 163.82, 162.95, 162.70, 159.19, 130.47, 129.26, 129.22, 129.08, 128.90, 115.14, 114.84, 114.26, 91.11, 91.04, 88.63, 56.06, 56.01, 55.88, 55.74, 55.40, 55.27, 43.56, 43.41, 42.87, 42.78, 40.43.

HRMS m/z calc. for $C_{22}H_{26}ClFN_3O_6S$ $[M+H]^+$ 514.1215 found 514.1212.

**Synthesis of alkyne-containing analogs of (*R,R*)-GL-373**

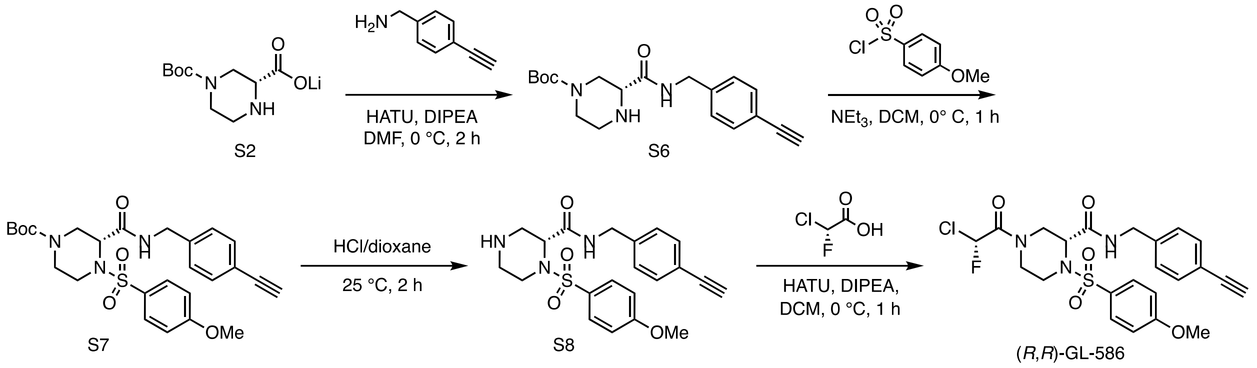


**Figure S6.** Synthesis of (*R*,*R*)-GL-586.

***tert*-butyl (*R*)-3-((4-ethynylbenzyl)carbamoyl)piperazine-1-carboxylate (S6)**

To a solution of **S2** (300 mg, 1.27 mmol, 1 equiv) and 4-ethynylbenzylamine (234 mg, 1.40 mmol, 1.1 equiv) in DMF (5 mL) were added DIPEA (492 mg, 3.81 mmol, 3.0 equiv) and HATU (724 mg, 1.91 mmol, 1.5 equiv). The mixture was stirred at 0 °C for 2 h. On completion, the reaction mixture was diluted with $H_2O$ (20 mL) and extracted with EtOAc (20 mL × 3). The combined organic layers were dried over $Na_2SO_4$, filtered and concentrated under reduced pressure to give a residue and purified by reversed-phase HPLC (A: $H_2O$ (10 mM $NH_4HCO_3$), B: acetonitrile; gradient: B: 30-50%, 20 min) to give **S6** (150 mg, 77% purity) as a yellow solid. Further purification by reversed-phase HPLC (A: $H_2O$ (10 mM formic acid), B: acetonitrile; gradient: B: 30-50%, 20 min) gave **S6•HCO₂H** (100 mg, 20% yield) as a white solid.

$^1$H NMR (400 MHz, $CDCl_3$) δ 8.12 (s, 1H, formate), 7.45 (d, *J* = 8.0 Hz, 2H), 7.35 – 7.26 (m, 1H), 7.21 (d, *J* = 7.9 Hz, 2H), 4.43 (d, *J* = 5.9 Hz, 2H), 4.07 (dd, *J* = 13.5, 3.7 Hz, 1H), 3.95 – 3.68 (m, 1H), 3.68 – 3.43 (m, 4H), 3.31 – 2.88 (m, 3H), 2.81 (ddd, *J* = 12.8, 9.7, 3.4 Hz, 1H), 1.45 (s, 9H).

LC-MS m/z calc. for $C_{15}H_{18}N_3O_3$ $[M\text{-}tBu+2H]^+$ 288.1 found 288.2.

***tert*-butyl (*R*)-3-((4-ethynylbenzyl)carbamoyl)-4-((4-methoxyphenyl)sulfonyl)piperazine-1-carboxylate (S7)**

To a precooled (0 °C) solution of **S6•HCO₂H** (100 mg, 0.257 mmol) in dichloromethane (5 mL) were added triethylamine (84.9 mg, 0.839 mmol, 3.3 equiv) and 4-methoxysulfonyl chloride (116 mg, 0.559 mmol, 2.2 equiv). The mixture was stirred at 0 °C for 1 h. On completion, the reaction mixture was concentrated under reduced pressure to give a residue, which was purified by flash silica gel chromatography ($SiO_2$, hexanes/EtOAc = 7:3 to 1:1) to give **S7** (40 mg, 26% yield) as a white solid.

$^1$H NMR (400 MHz, $CDCl_3$) δ 7.76 (d, *J* = 8.9 Hz, 2H), 7.45 (d, *J* = 8.2 Hz, 2H), 7.19 (d, *J* = 7.9 Hz, 2H), 6.98 (d, *J* = 8.9 Hz, 2H), 6.95 – 6.90 (m, 1H), 4.65 – 4.34 (m, 4H), 3.88 (s, 3H), 3.84 – 3.59 (m, 2H), 3.33 – 3.21 (m, 1H), 3.07 (s, 1H), 2.85 – 2.61 (m, 2H), 1.41 (s, 9H).

LC-MS m/z calc. for $C_{27}H_{32}N_3O_6S$ $[M+H]^+$ 514.2 found 514.1.

**(*R*)-*N*-(4-ethynylbenzyl)-1-((4-methoxyphenyl)sulfonyl)piperazine-2-carboxamide (S8)**

To a solution of **S7** (200 mg, 389 µmol) in dioxane (1 mL) was added HCl/dioxane (4 M, 2 mL) and the resulting mixture was stirred at 25 °C for 1 h. On completion, the reaction mixture was concentrated to give compound **S8•HCl** (160 mg, quant.) as a yellow solid, which was used in the next step without further purification.

**(*R*)-4-((*R*)-2-chloro-2-fluoroacetyl)-*N*-(4-ethynylbenzyl)-1-((4-methoxyphenyl)sulfonyl) piperazine-2-carboxamide ((*R,R*)-GL-586)**

To a precooled (0 ºC) solution of (*R*)-chlorofluoroacetic acid (26.1 mg, 232 µmol), HATU (147 mg, 387 µmol) and DIEA (75.0 mg, 580 µmol) in dichloromethane (10 mL) was added **S8•HCl** (80.0 mg, 193 µmol) and the mixture was stirred at 0 °C for 2 h. On completion, the reaction mixture was partitioned between ethyl acetate (60 mL) and brine (40 mL). Then, the aqueous layer was extracted with ethyl acetate (40 mL x 3). The organic layers were combined, dried over sodium sulfate, filtered and concentrated under reduced pressure to give a residue, which was purified by prep-TLC ($SiO_2$, petroleum ether/EtOAc = 2:1) and prep-HPLC (column: Waters Xbridge 150*25mm* 5um; mobile phase: A: $H_2O$ (10 mM $NH_4HCO_3$), B: acetonitrile; gradient: B: 40%-70% over 9 min) to give **(*R,R*)-GL-586** (26.0 mg, 26% yield) as a white solid.

Note: NMR spectra are consistent with a 3:2 mixture of rotamers.

$^1$H NMR (400 MHz, $CD_3OD$) δ 7.82 – 7.69 (m, 2H), 7.42 (d, *J* = 8.2 Hz, 2H), 7.23 (d, *J* = 7.9 Hz, 2H), 7.01 (t, *J* = 9.6 Hz, 2H), 6.91 (d, $^2J_{HF}$ = 49.2 Hz, 0.6H), 6.84 (d, $^2J_{HF}$ = 49.1 Hz, 0.4H), 4.70 (d, *J* = 13.7 Hz, 0.6H), 4.61 – 4.48 (m, 1H), 4.40 – 4.16 (m, 2.4H), 4.08 (br d, *J* = 13.7 Hz, 0.4H), 3.94 (d, *J* = 13.8 Hz, 0.6H), 3.87 (s, 3H), 3.83 – 3.71 (m, 1.6H), 3.70 – 3.58 (m, 0.4H), 3.46 (s, 1H), 3.39 (dd, *J* = 14.1, 4.7 Hz, 0.4H), 3.24 (ddd, *J* = 14.6, 9.9, 5.6 Hz, 0.6H), 3.09 (dd, *J* = 13.7, 4.7 Hz, 0.6H), 3.00 (t, *J* = 13.0 Hz, 0.4H), 1 exchangeable proton not observed.

HRMS m/z calc. for $C_{23}H_{24}ClFN_3O_5S$ $[M+H]^+$ 508.1109 found 508.1108.

**(*S*)-4-((*S*)-2-chloro-2-fluoroacetyl)-*N*-(4-ethynylbenzyl)-1-((4-methoxyphenyl)sulfonyl) piperazine-2-carboxamide ((*S,S*)-GL-586)**

**(*S,S*)-GL-586** was prepared from *ent*-**S8•HCl** and (*S*)-chlorofluoroacetic acid following a procedure analogous to that used for the synthesis of (*R*,*R*)-GL-586.

Note: NMR spectra are consistent with a 3:2 mixture of rotamers.

$^1$H NMR (400 MHz, MeOD) δ 7.81 – 7.70 (m, 2H), 7.42 (d, *J* = 8.0 Hz, 2H), 7.23 (d, *J* = 7.9 Hz, 2H), 7.01 (t, *J* = 9.8 Hz, 2H), 6.91 (d, $^2J_{HF}$ = 49.2 Hz, 0.6H), 6.84 (d, $^2J_{HF}$ = 49.1 Hz, 0.4H), 4.70 (d, *J* = 13.7 Hz, 0.6H), 4.58 – 4.50 (m, 1H), 4.40 – 4.16 (m, 2.4H), 4.12 – 4.04 (m, 0.4H), 3.95 (br d, *J* = 13.8 Hz, 0.6H), 3.87 (s, 3H), 3.84 – 3.71 (m, 1.6H), 3.70 – 3.57 (m, 0.4H), 3.46 (s, 1H), 3.39 (dd, *J* = 14.0, 4.6 Hz, 0.4H), 3.24 (ddd, *J* = 14.7, 10.0, 5.6 Hz, 0.6H), 3.09 (dd, *J* = 13.7, 4.7 Hz, 0.6H), 3.05 – 2.91 (m, 0.4H), 1 exchangeable proton not observed.

HRMS m/z calc. for $C_{23}H_{24}ClFN_3O_5S$ $[M+H]^+$ 508.1109 found 508.1118.

### Analytical data: NMR spectra

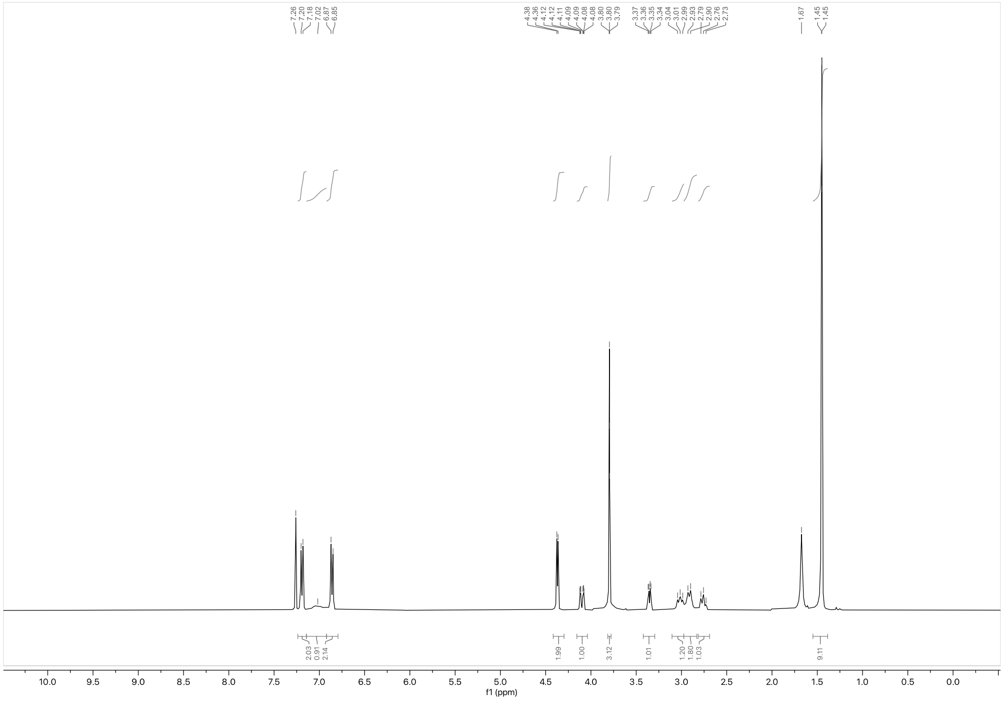

$^1$H NMR spectrum of **S3** (400 MHz, $CDCl_3$)

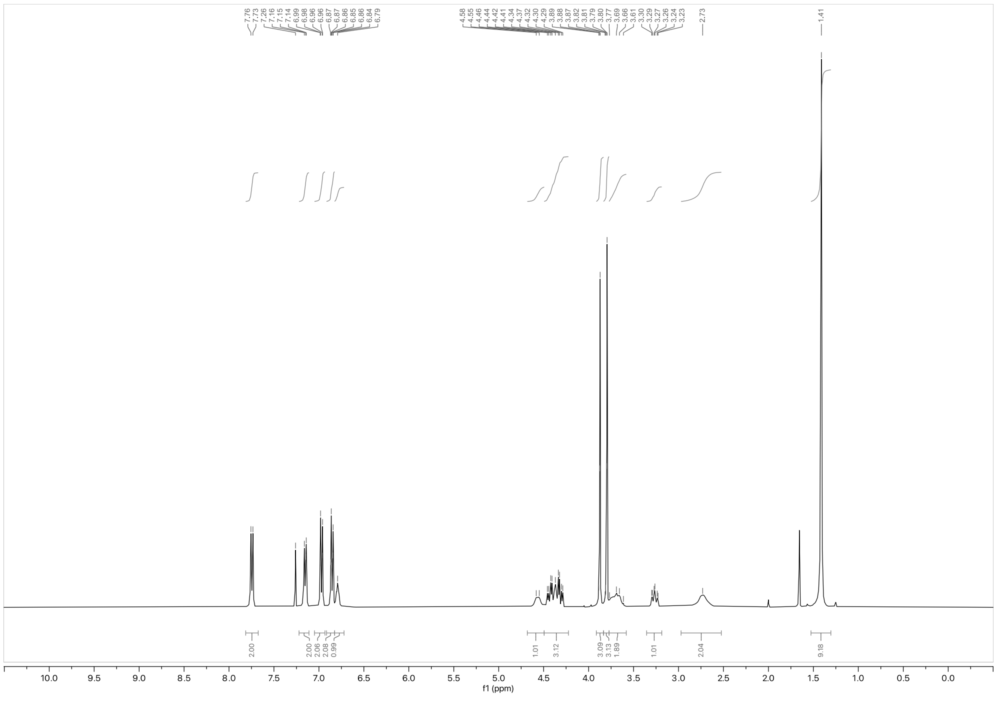

[1]H NMR spectrum of **S4** (400 MHz, $CDCl_3$)

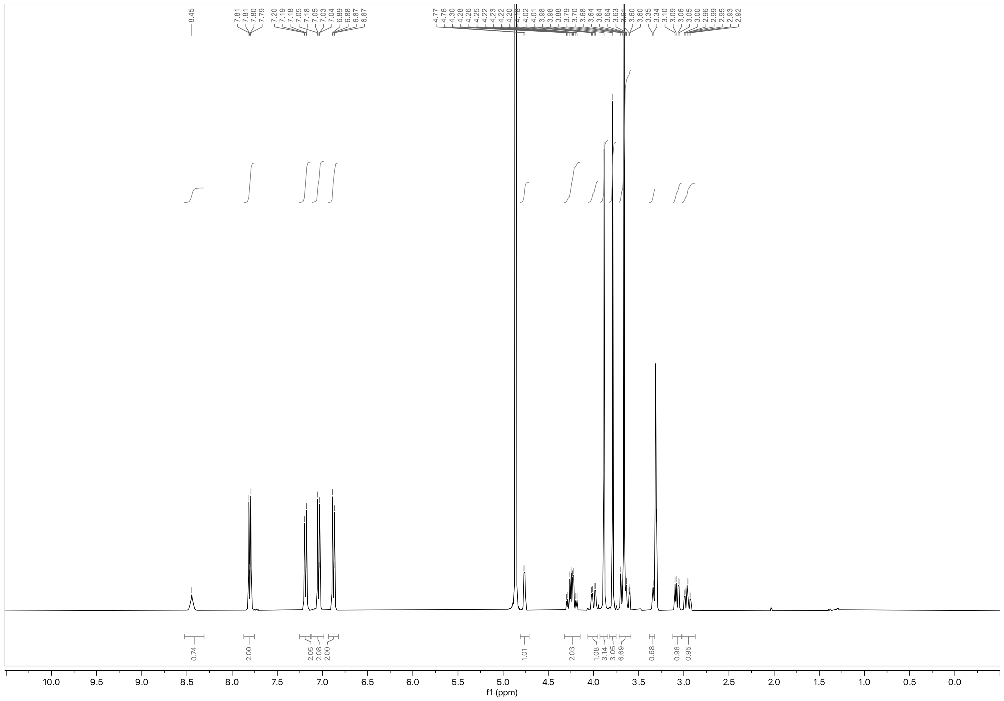

$^1$H NMR spectrum of **S5•HCl** (400 MHz, $CD_3OD$)

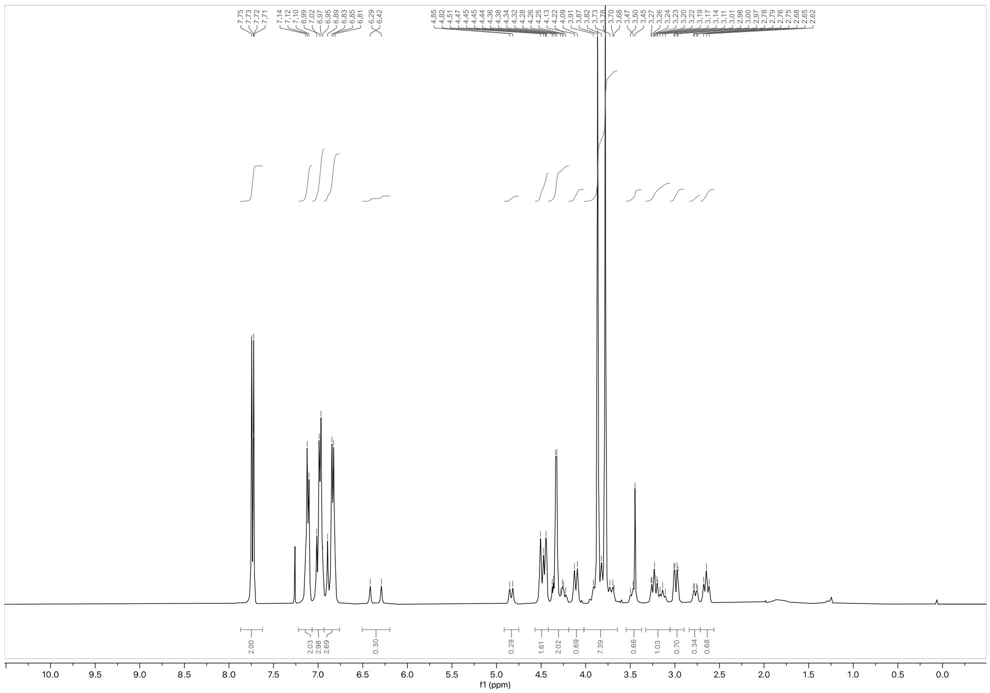

$^1$H NMR spectrum of **(*R*,*R*)-GL-373** (400 MHz, $CDCl_3$)

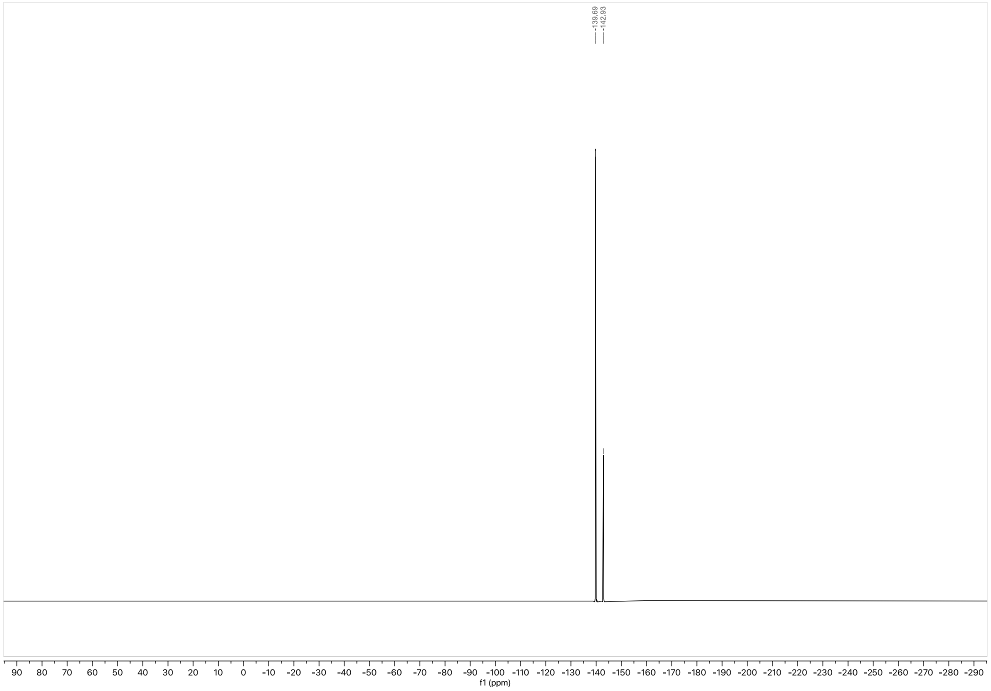


$^{19}F$ NMR spectrum of ***(R,R)*-GL-373** (376 MHz, $CDCl_3$)

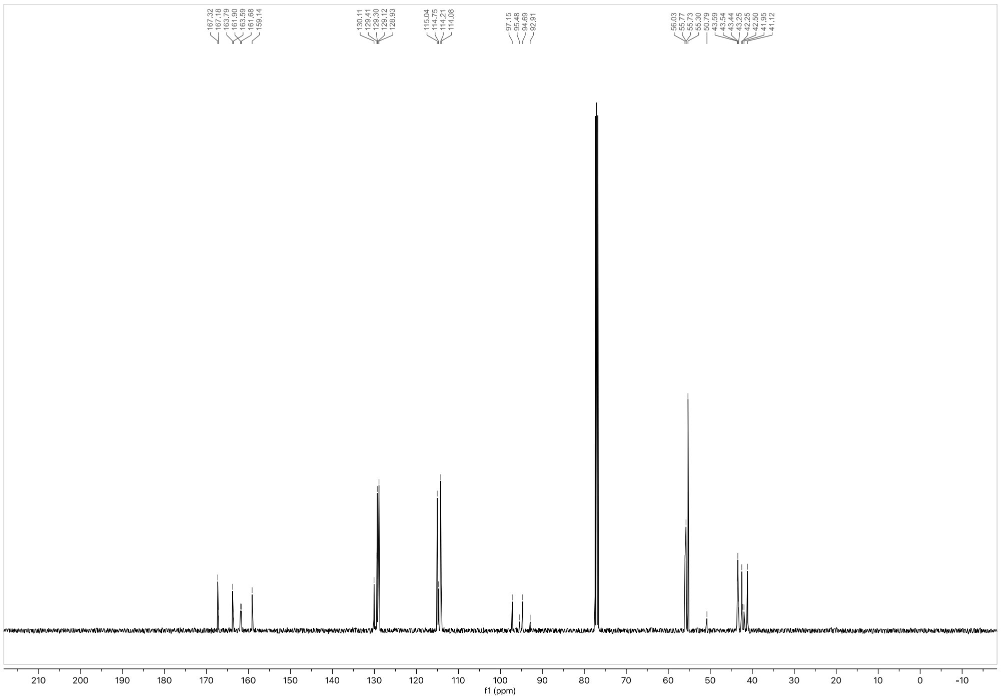


$^{13}C$ NMR spectrum of ***(R,R)*-GL-373** (101 MHz, $CDCl_3$)

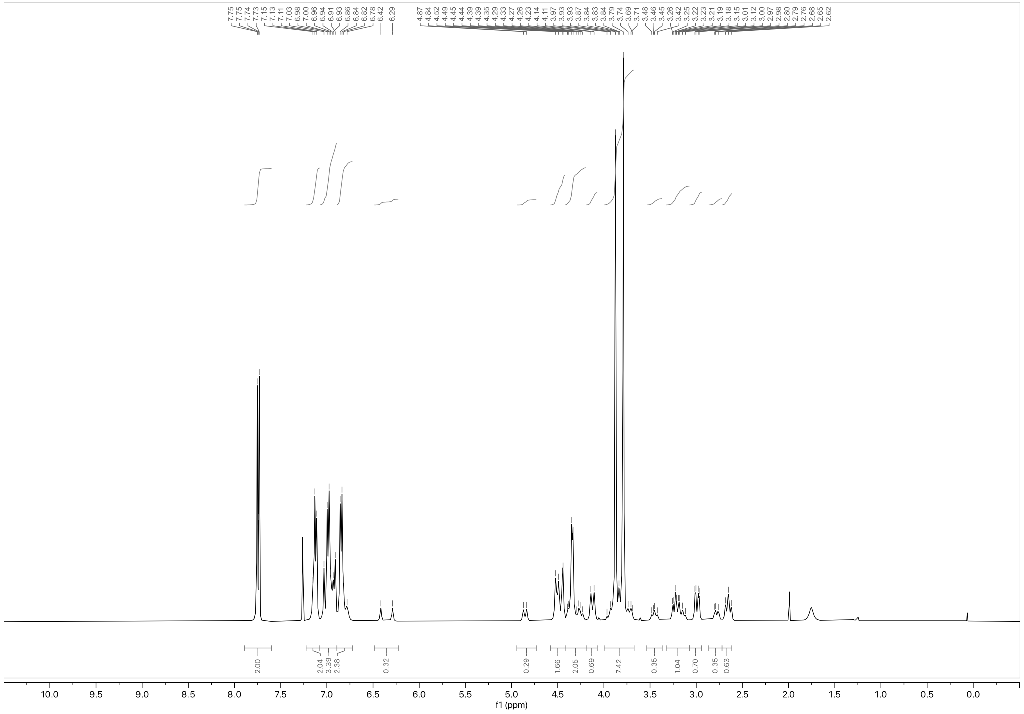

$^{1}$H NMR spectrum of **(*S*,*S*)-GL-373** (400 MHz, $CDCl_3$)

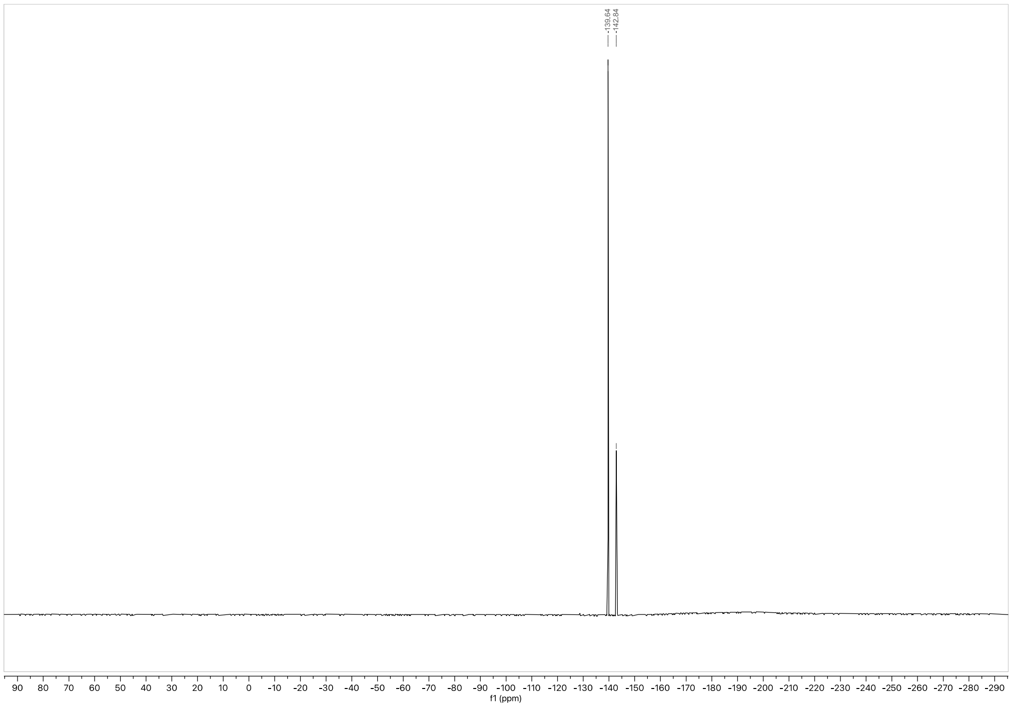

$^{19}$F NMR spectrum of **(*S*,*S*)-GL-373** (376 MHz, $CDCl_3$)

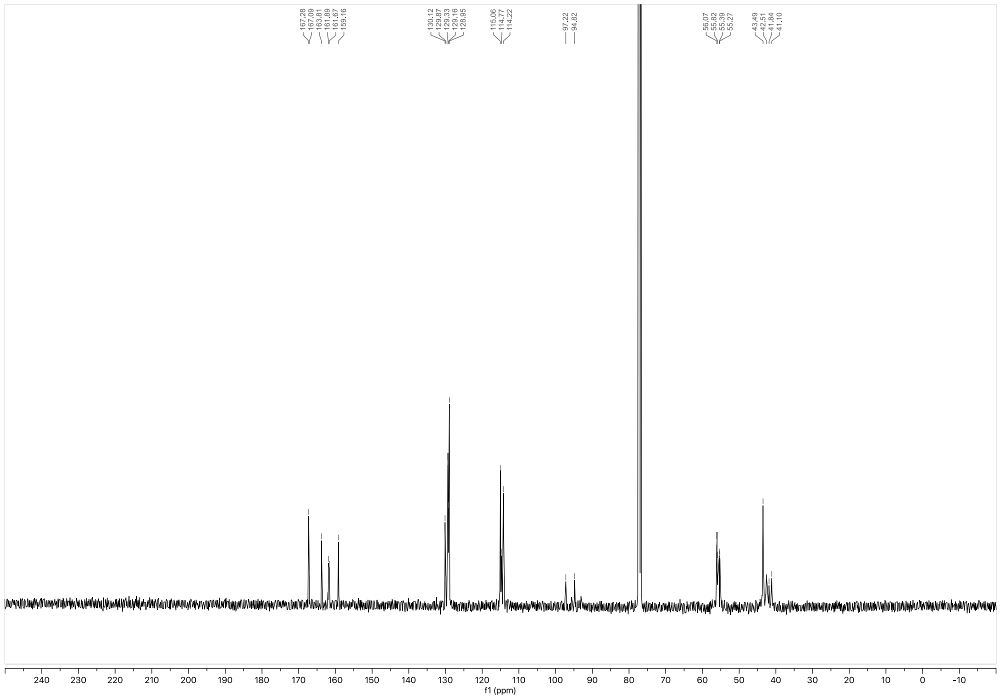

$^{13}C$ NMR spectrum of ***(S,S)*-GL-373** (101 MHz, $CDCl_3$)

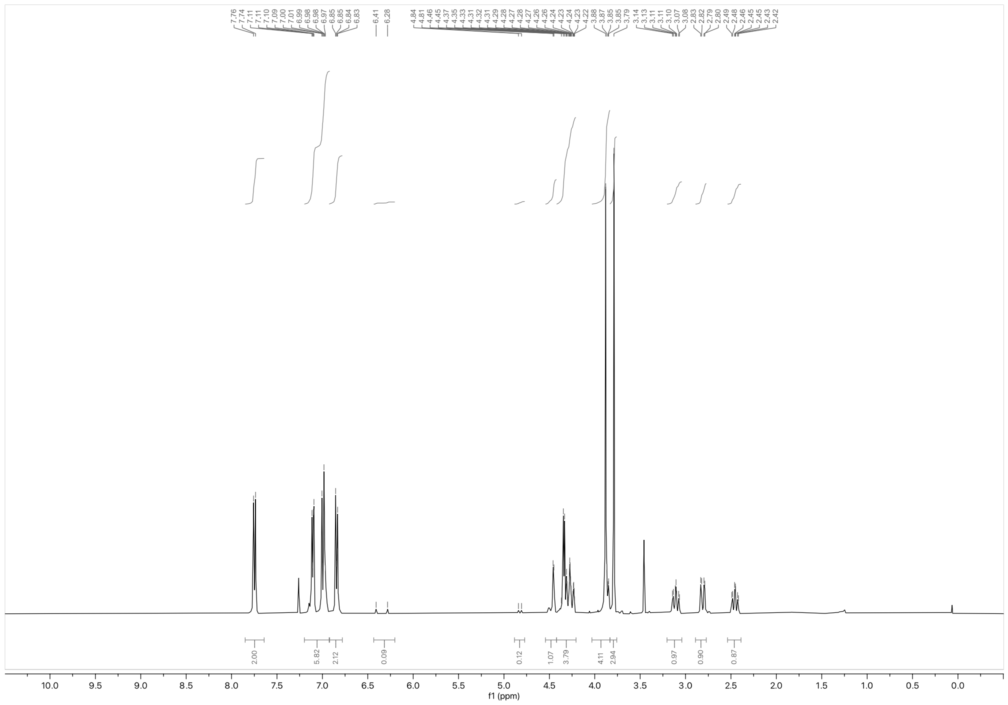

$^{1}H$ NMR spectrum of ***(S,R)*-GL-373** (400 MHz, $CDCl_3$)

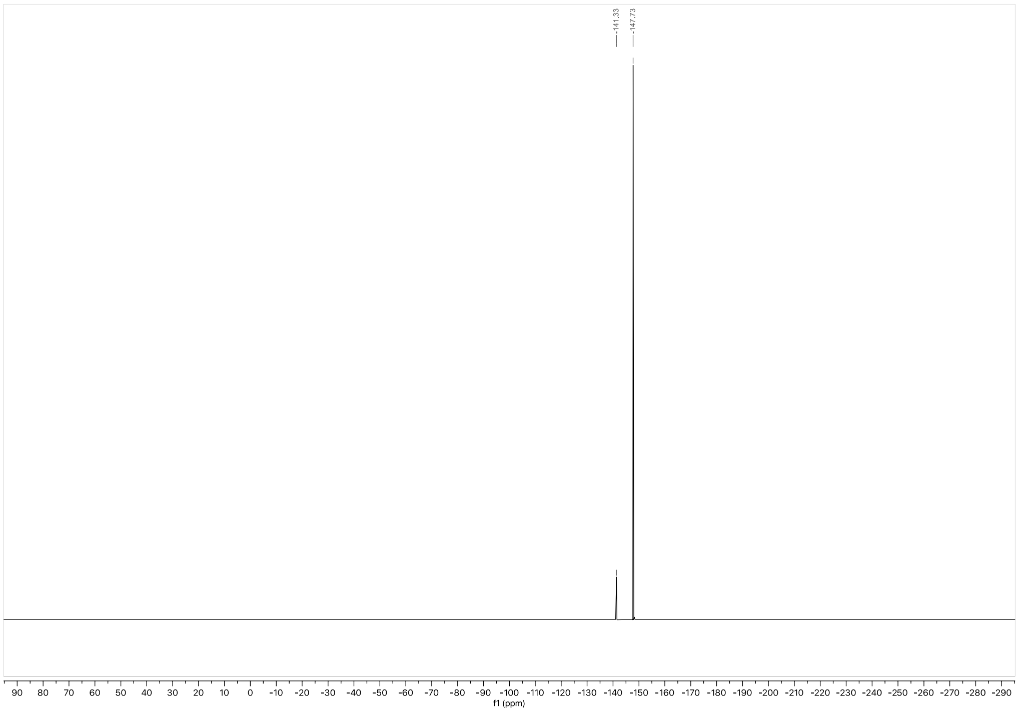


$^{19}$F NMR spectrum of **(*S*,*R*)-GL-373** (376 MHz, $CDCl_3$)

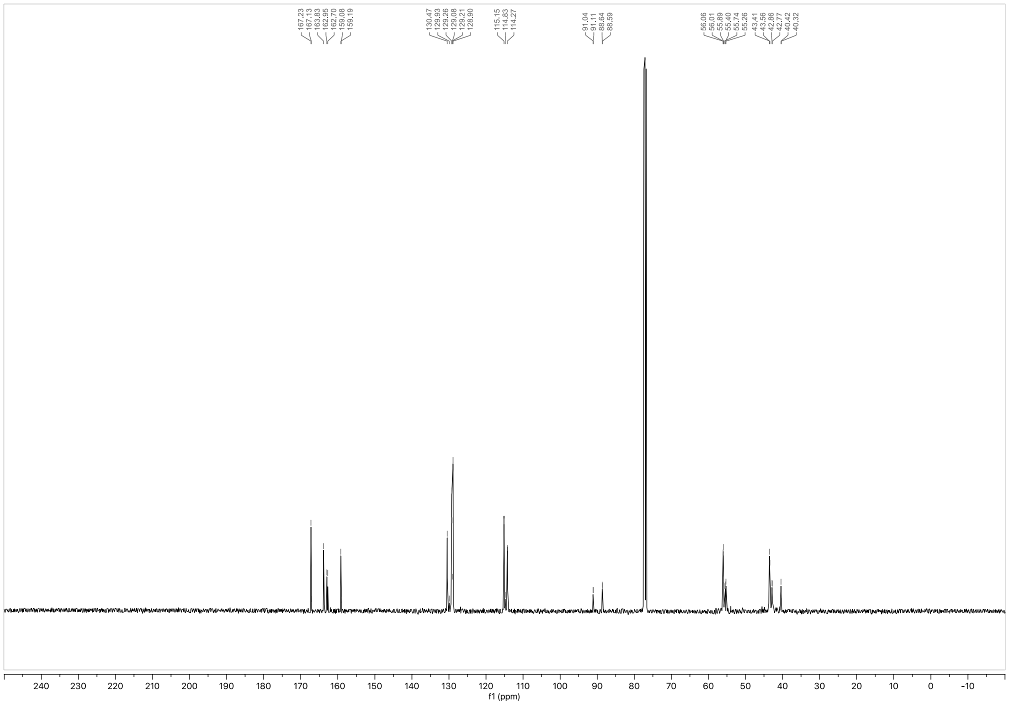


$^{13}$C NMR spectrum of **(*S*,*R*)-GL-373** (101 MHz, $CDCl_3$)

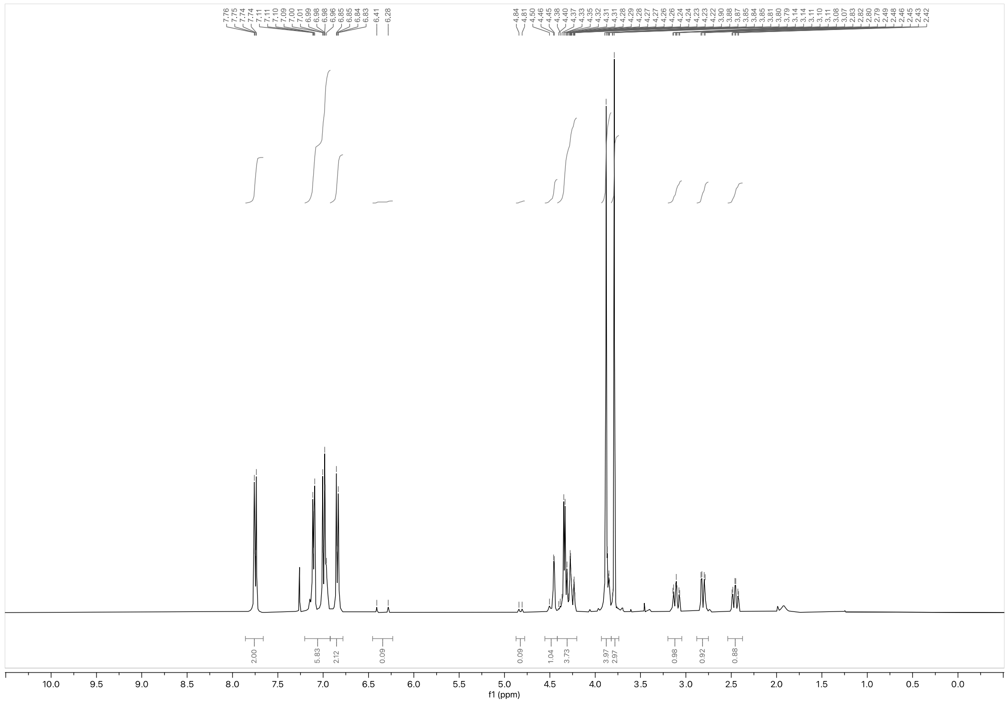

$^{1}H$ NMR spectrum of ***(R,S)*-GL-373** (400 MHz, $CDCl_3$)

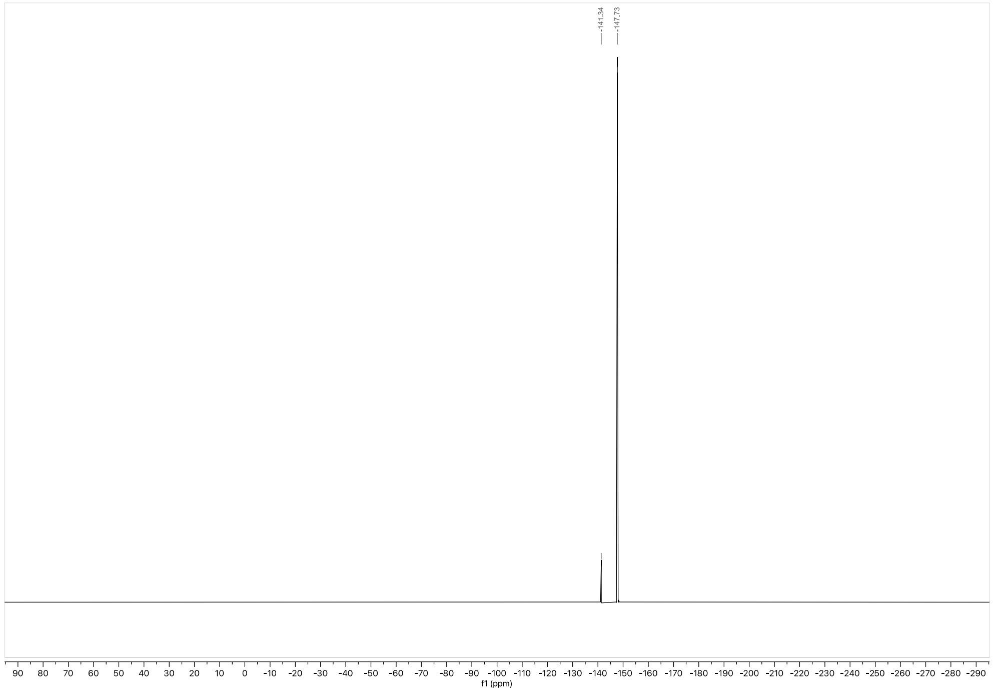

$^{19}F$ NMR spectrum of ***(R,S)*-GL-373** (376 MHz, $CDCl_3$)

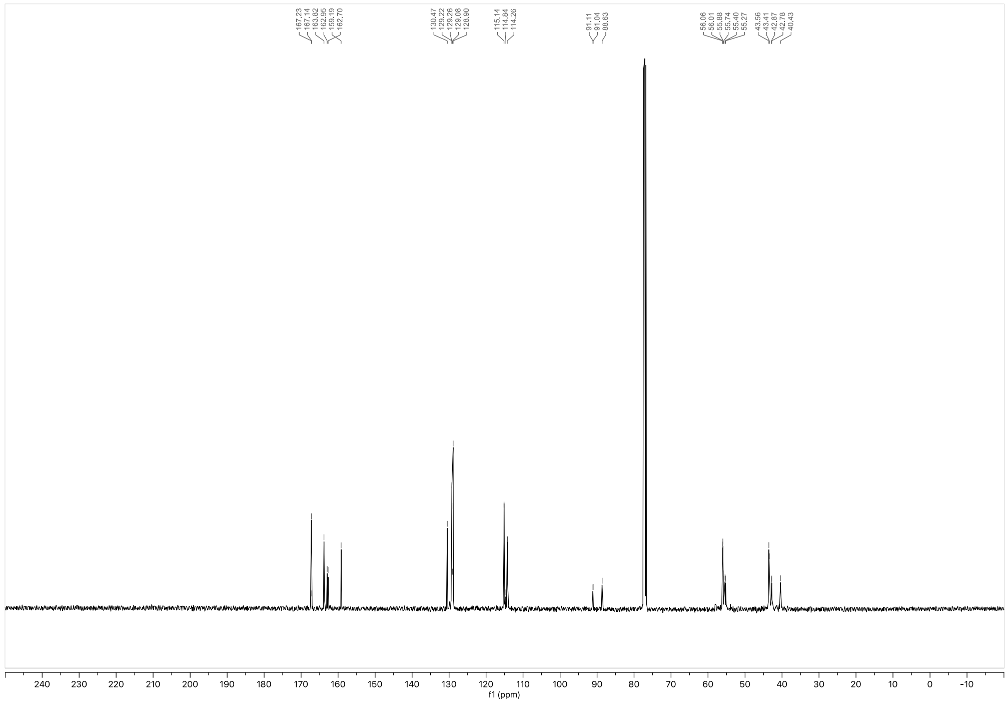

[13] C NMR spectrum of ***(R,S)*-GL-373** (101 MHz, $CDCl_3$)

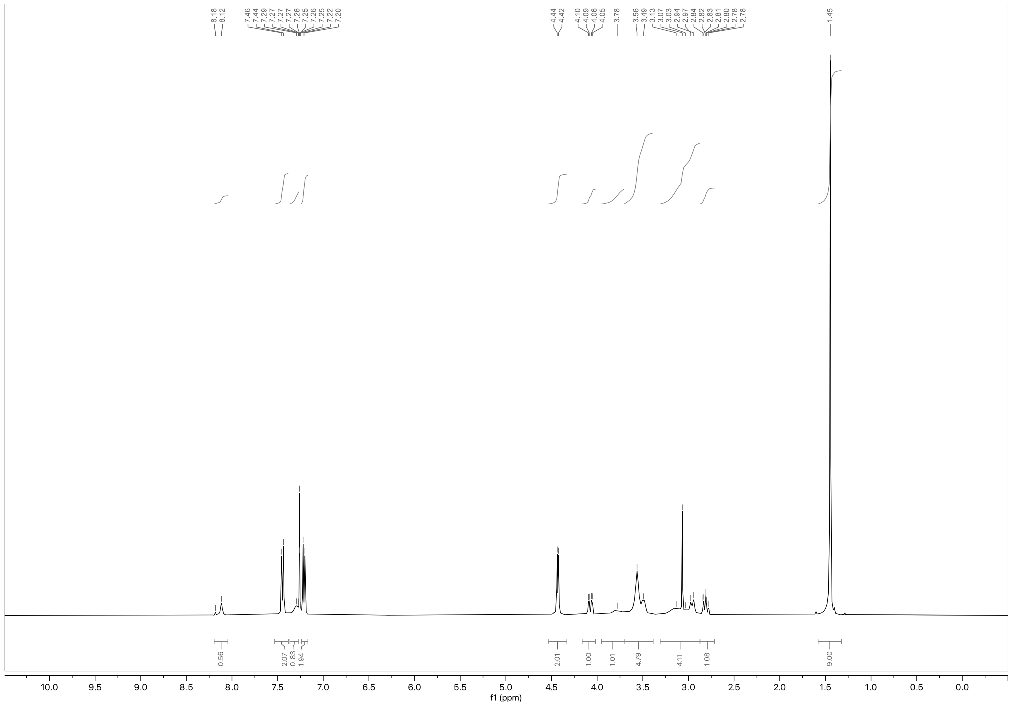

[1] H NMR spectrum of **S6•HCO₂H** (400 MHz, $CDCl_3$)

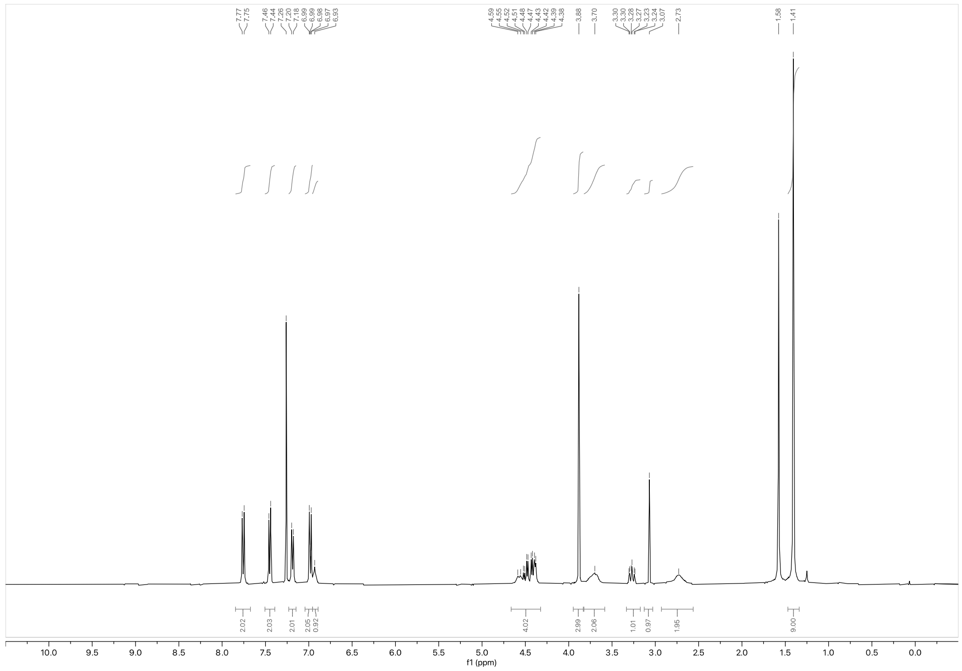

[1]H NMR spectrum of **S7** (400 MHz, $CDCl_3$)

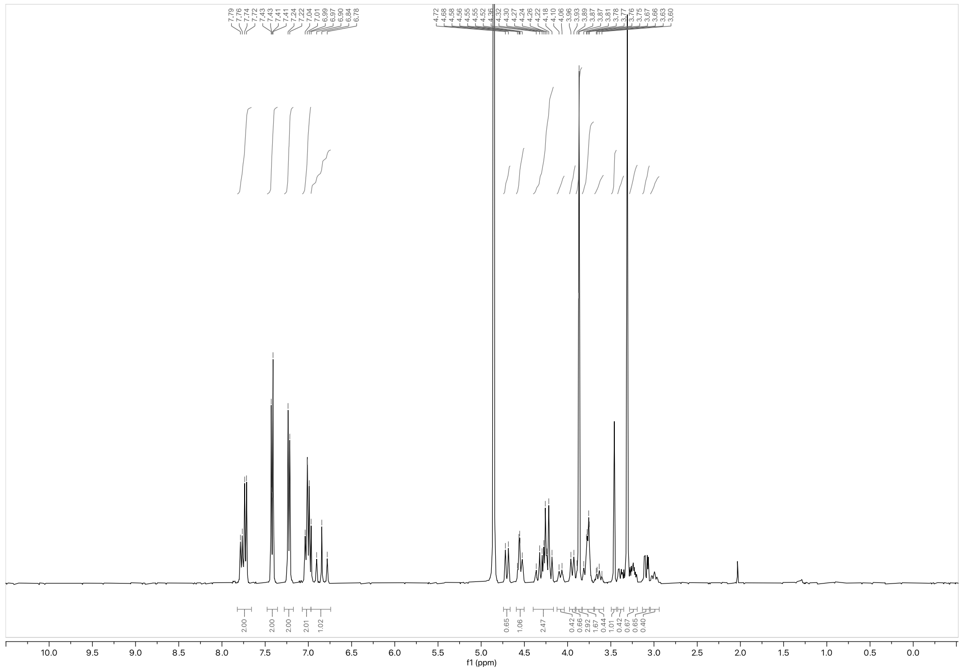

[1]H NMR spectrum of ***(R,R)*-GL-586** (400 MHz, $CD_3OD$)

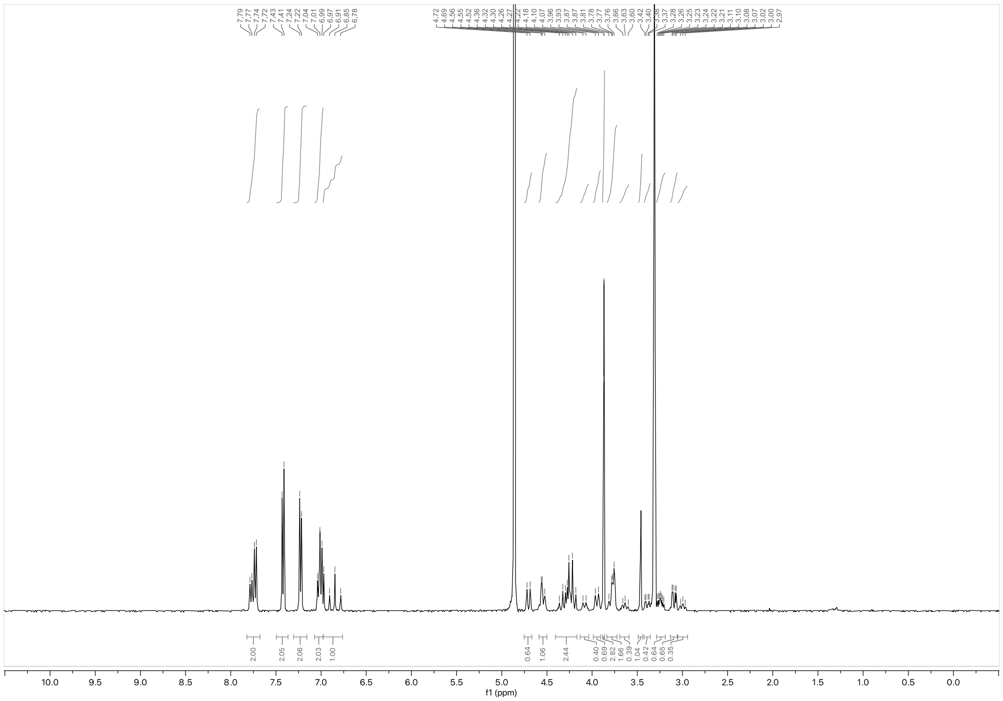

$^{1}H$ NMR spectrum of **(*S*,*S*)-GL-586** (400 MHz, $CD_3OD$)

**Analytical data: SFC**

(*R*,*R*)-GL-373

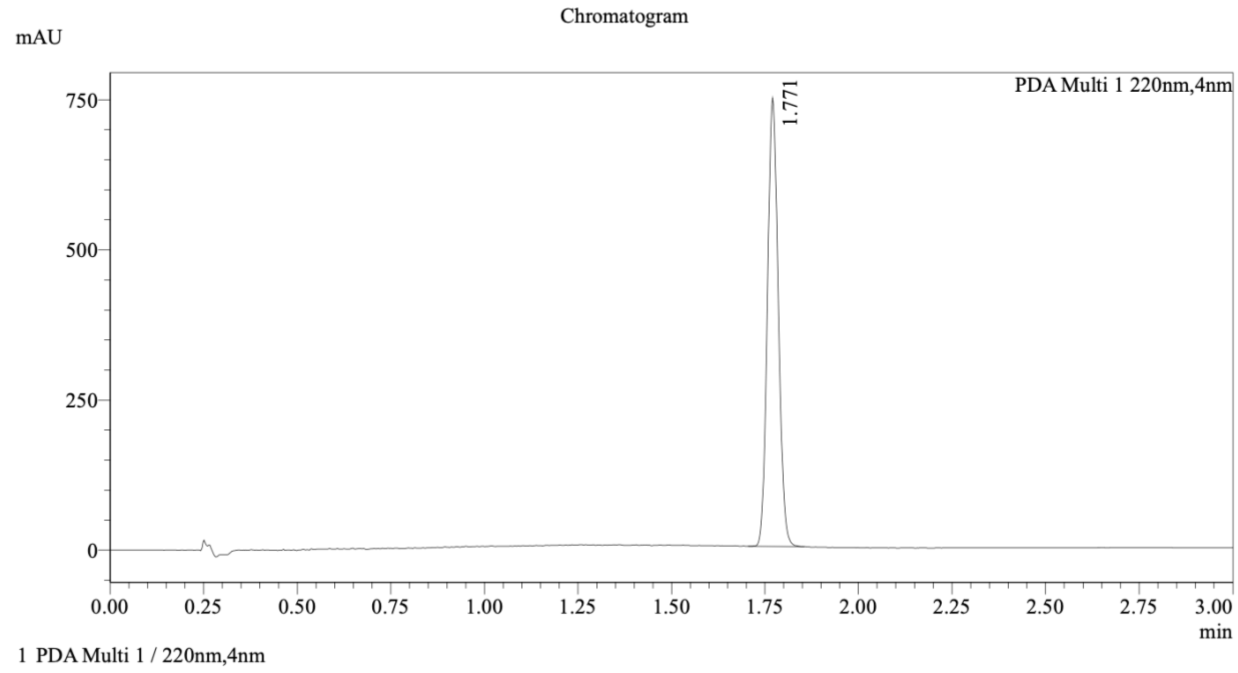


(*S*,*S*)-GL-373

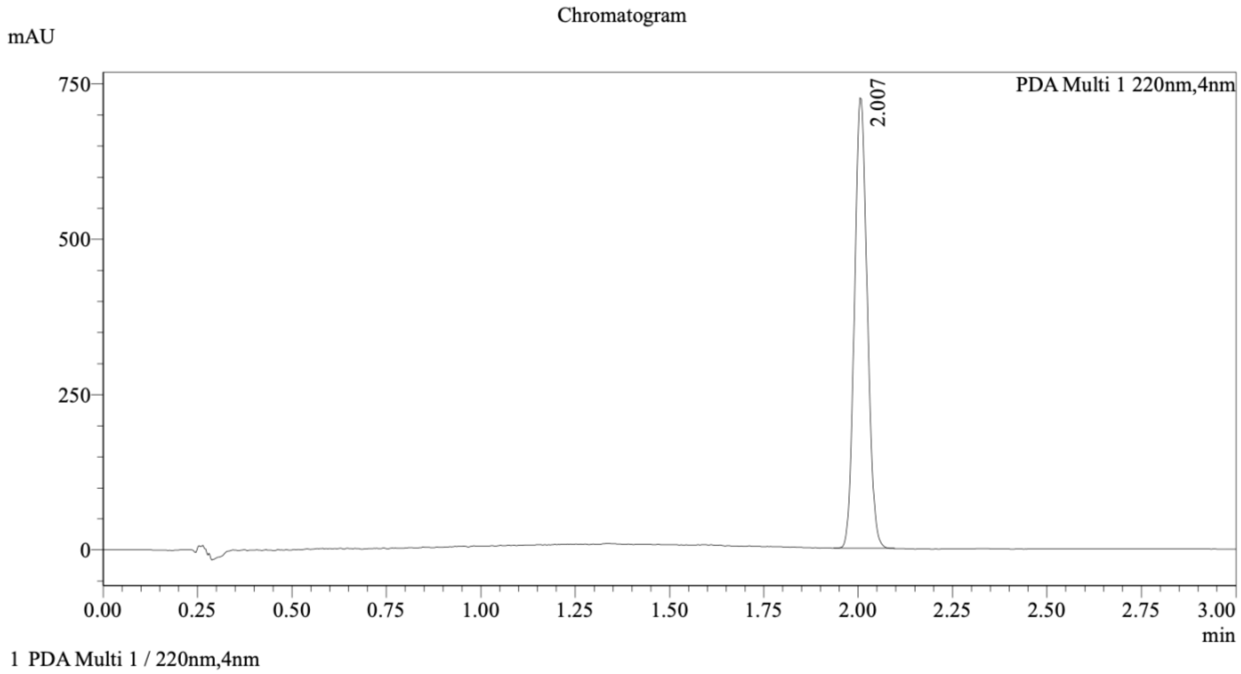

Mixture of (*R*,*R*)-GL-373 and (*S*,*S*)-GL-373

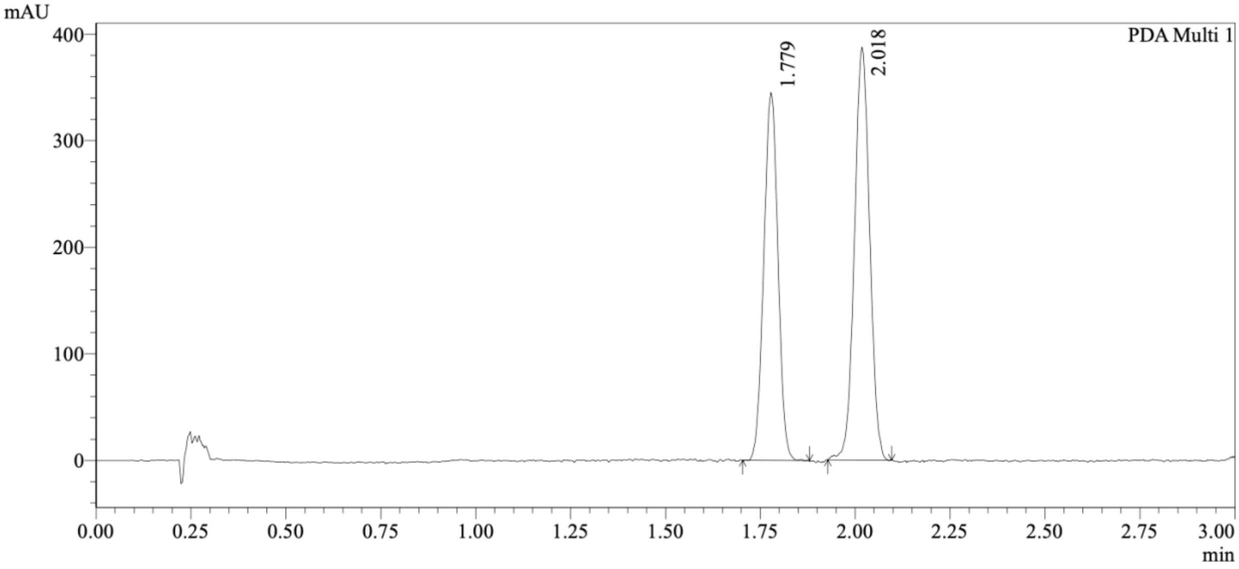


1 PDA Multi 1 / 220nm,4nm

Integration Results

PeakTable

PDA Ch1 220nm

| Peak# | Ret. Time | USP Width | Resolution | Height | Area | Area % |
|---|---|---|---|---|---|---|
| 1 | 1.779 | 0.072 | 0.000 | 339037 | 894991 | 45.270 |
| 2 | 2.018 | 0.077 | 3.223 | 378899 | 1081996 | 54.730 |
| Total | | | | 717936 | 1976987 | 100.000 |

Method Details

Column: Chiralcel OJ-3 50×4.6mm I.D., 3 µm

Mobile phase: A: $CO_2$, B: MeOH (0.05% diethylamine)

Gradient elution: 5% to 40% B

(*S*,*R*)-GL-373

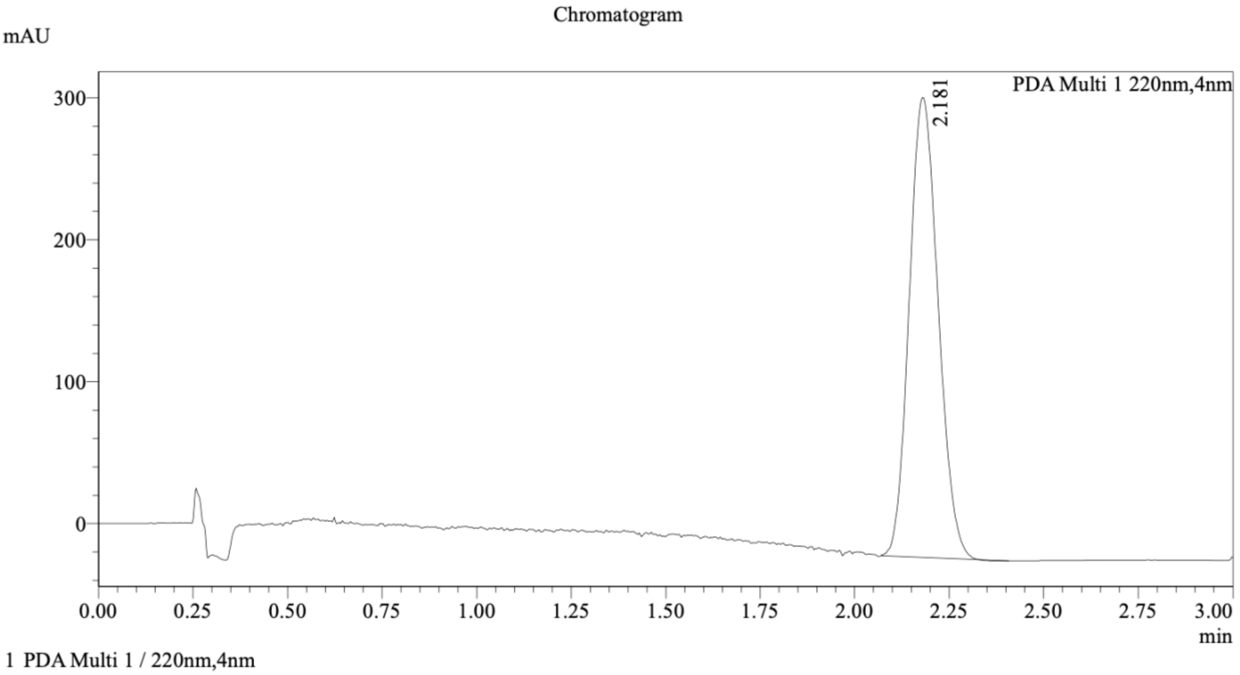


(*R*,*S*)-GL-373

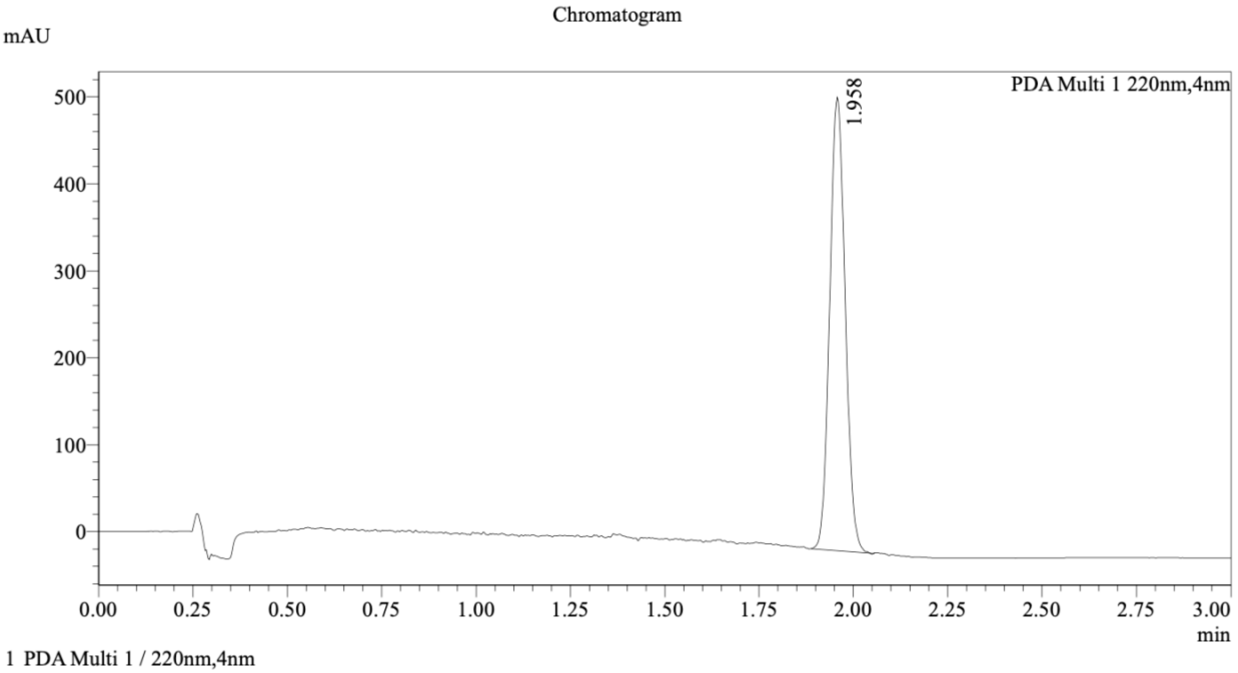

Mixture of (*S*,*R*)-GL-373 and (*R*,*S*)-GL-373

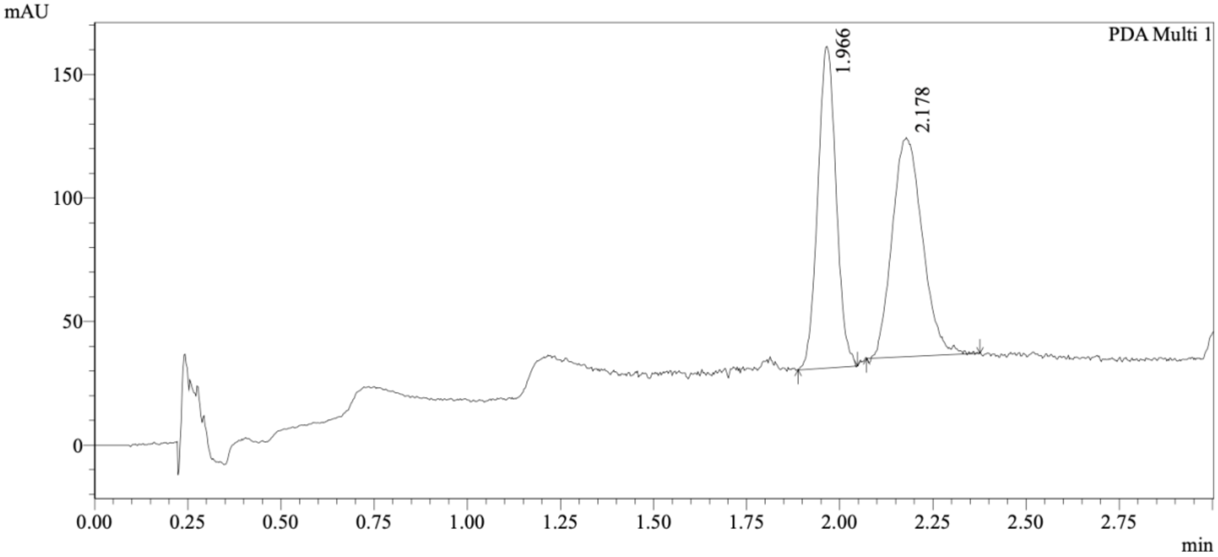


1 PDA Multi 1 / 220nm,4nm

Integration Results

PeakTable

PDA Ch1 220nm

| Peak# | Ret. Time | USP Width | Resolution | Height | Area | Area % |
|---|---|---|---|---|---|---|
| 1 | 1.966 | 0.092 | 0.000 | 129725 | 447477 | 47.237 |
| 2 | 2.178 | 0.150 | 1.755 | 87979 | 499834 | 52.763 |
| Total | | | | 217704 | 947311 | 100.000 |

Method Details

Column: Chiralcel OJ-3 50×4.6mm I.D., 3 μm

Mobile phase: A: $CO_2$, B: EtOH (0.05% diethylamine)

Gradient elution: 5% to 40% B

(*R*,*R*)-GL-586

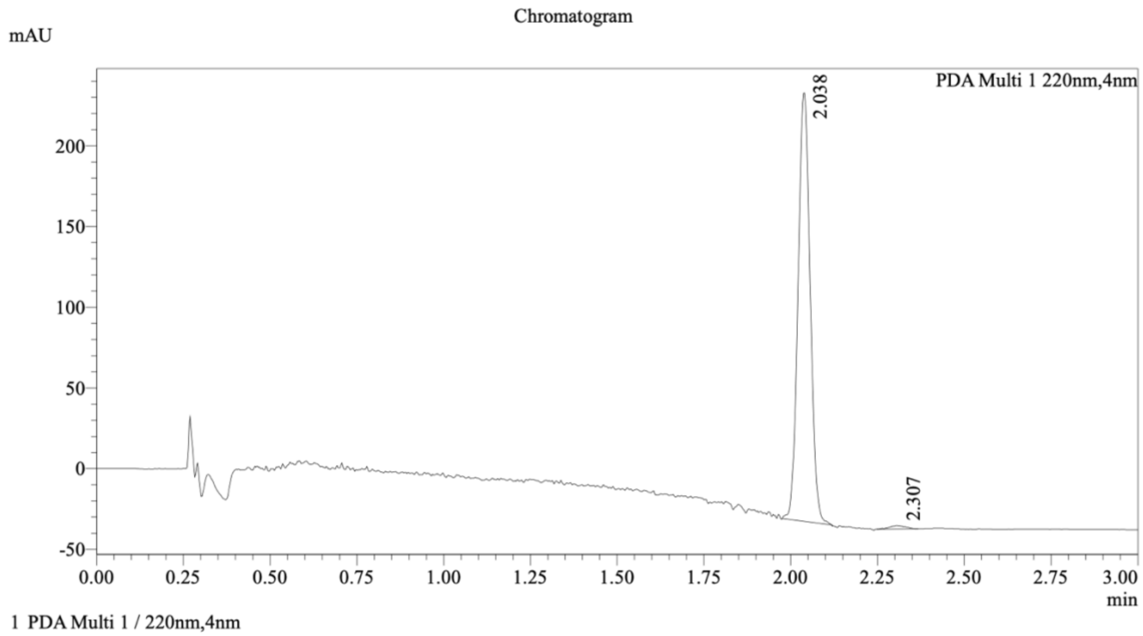


Integration Result

Peak Table

PDA Ch1 220nm

| Peak# | Ret. Time | Height | Height% | Resolution(USP) | Area | Area% |
|---|---|---|---|---|---|---|
| 1 | 2.038 | 263042 | 99.249 | -- | 644125 | 98.932 |
| 2 | 2.307 | 1990 | 0.751 | 3.273 | 6956 | 1.068 |

(*S*,*S*)-GL-586

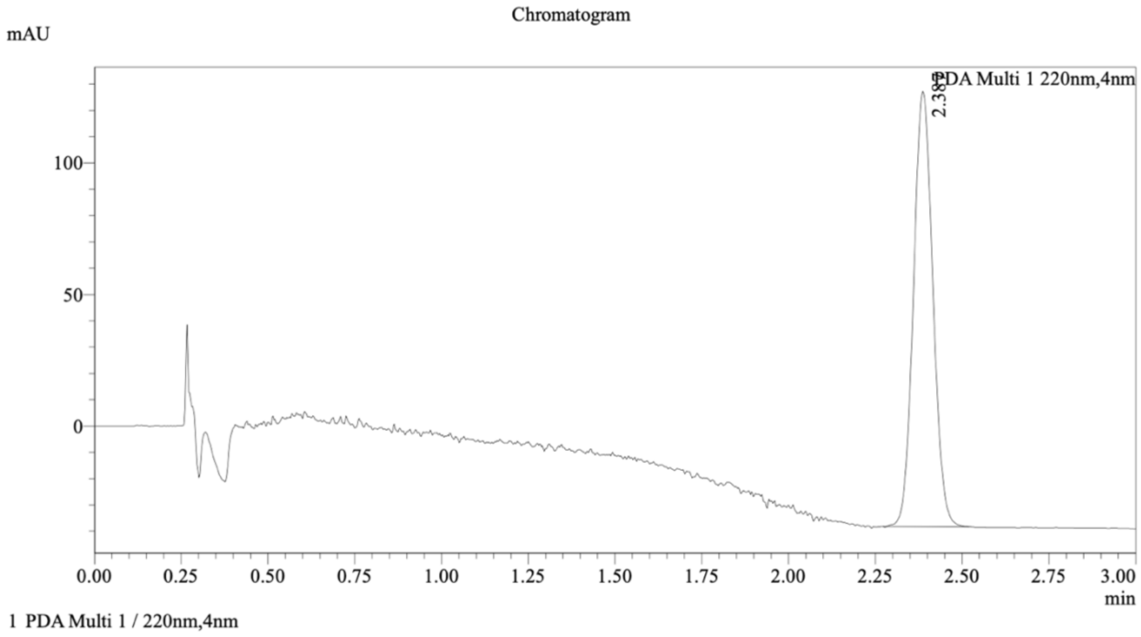
Chromatogram
mAU
PDA Multi 1 220nm,4nm
2.389
100
50
0
0.00
0.25
0.50
0.75
1.00
1.25
1.50
1.75
2.00
2.25
2.50
2.75
3.00
min
1 PDA Multi 1 / 220nm,4nm

Mixture of (*R*,*R*)-GL-586 and (*S*,*S*)-GL-586

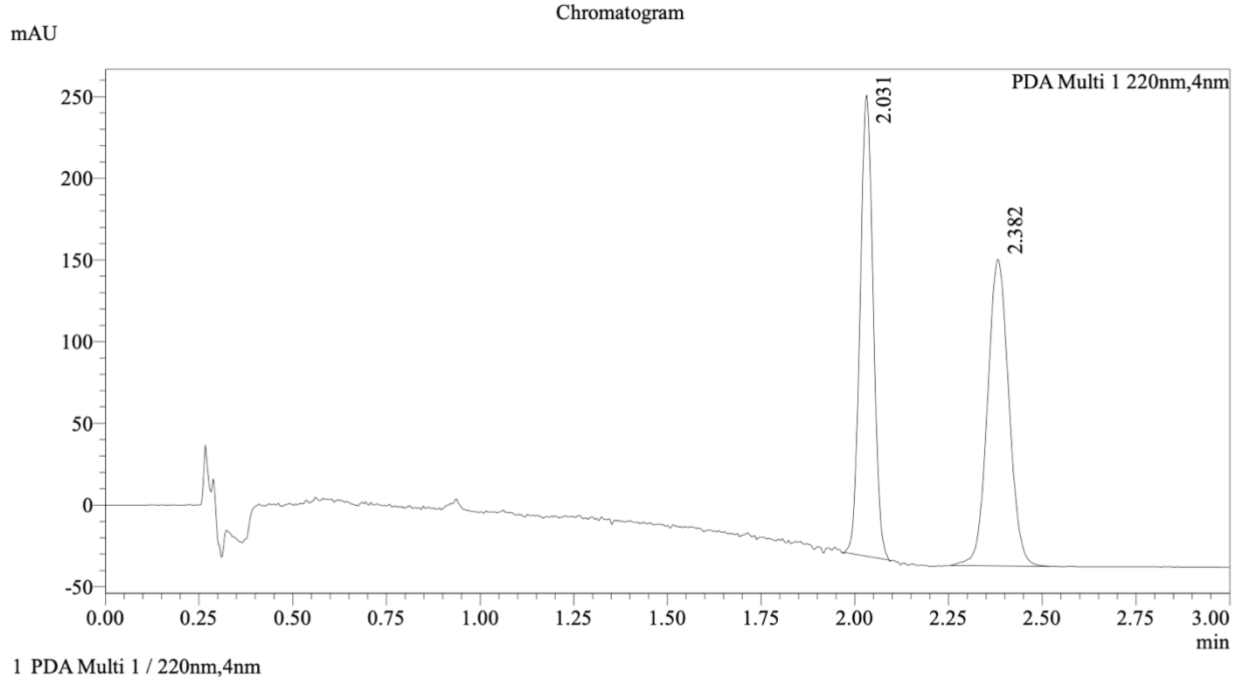


Integration Result

Peak Table

PDA Ch1 220nm

| Peak# | Ret. Time | Height | Height% | Resolution(USP) | Area | Area% |
|---|---|---|---|---|---|---|
| 1 | 2.031 | 277506 | 59.747 | -- | 688514 | 48.048 |
| 2 | 2.382 | 186960 | 40.253 | 4.021 | 744468 | 51.952 |

Method Details

Column: Chiralcel OJ-3 50×4.6mm I.D., 3 μm

Mobile phase: A: $CO_2$, B: EtOH (0.05% diethylamine)

Gradient elution: 5% to 40% B

Organics, and Gases in Deuterated Solvents Relevant to the Organometallic Chemist. Organometallics *29*, 2176-2179. 10.1021/om100106e.
